# Semantic integration demands modulate large-scale network interactions in the brain

**DOI:** 10.1101/2024.11.20.624508

**Authors:** Laura Nieberlein, Sandra Martin, Kathleen A. Williams, Alexander Gussew, Sophia D. Cyriaks, Maximilian Scheer, Stefan Rampp, Julian Prell, Gesa Hartwigsen

## Abstract

The ability to integrate semantic information into the context of a sentence is essential for human communication. Several studies have shown that the predictability of a final keyword based on the sentence context influences semantic integration on the behavioral, neurophysiological, and neural level. However, the architecture of the underlying network interactions for semantic integration across the lifespan remains unclear.

In this study, 32 healthy participants (30-75 years) performed an auditory cloze probability task during functional magnetic resonance imaging (fMRI), requiring lexical decisions on the sentence’s final words. Semantic integration demands were implicitly modulated by presenting sentences with expected, unexpected, anomalous, or pseudoword endings. To elucidate network interactions supporting semantic integration, we combined univariate task-based fMRI analyses with seed-based connectivity and between-network connectivity analyses.

Behavioral data revealed typical semantic integration effects, with increased integration demands being associated with longer response latencies and reduced accuracy. Univariate results demonstrated increased left frontal and temporal brain activity for sentences with higher integration demands. Between-network interactions highlighted the role of task-positive and default mode networks for sentence processing with increased semantic integration demands. Furthermore, increasing integration demands led to a higher number of behaviorally relevant network interactions, suggesting that the increased between-network coupling becomes more relevant for successful task performance as integration demands increase.

Our findings elucidate the complex network interactions underlying semantic integration across the aging continuum. Stronger interactions between various task-positive and default mode networks correlated with more efficient processing of sentences with increased semantic integration demands. These results may inform future studies with healthy old and clinical populations.

## Introduction

Efficient language comprehension in everyday communication requires the integration of context-dependent information into sentence meaning, a process known as semantic integration. There is consensus that the combination of words into a sentence generates expectations about further information (e.g., Kutas & Hillyard, 1980), and the human brain constantly generates context-dependent predictions about the subsequent input (for a review, see Ryskin & Nieuwland, 2023). Numerous studies have shown that the integration of a final keyword into a sentence is influenced by the preceding context (e.g., Hagoort, 2006; Hagoort & Indefrey, 2014; Obleser & Kotz, 2010; Rogalsky & Hickok, 2009; Zhu et al., 2013). However, expectations about subsequent input are frequently violated in everyday communication, for example, when the less dominant meaning of an ambiguous word is processed or when we perceive jokes or ironic statements. Such situations require additional integration processes during lexical access (Franzmeier et al., 2012).

At the behavioral level, higher semantic integration demands are reflected in increased response latencies and error rates (e.g., Baumgaertner et al., 2002; Kutas & Federmeier, 2000; Kutas & Hillyard, 1980; Lau et al., 2008). At the neurophysiological or neural level, integration demands result in increased N400 amplitudes (DeLong et al., 2014; Federmeier, 2007; Kutas & Federmeier, 2000; Kutas & Hillyard, 1980; Lau et al., 2008) and increased task-related activity in frontal and temporal brain areas (Baumgaertner et al., 2002; Hartwigsen et al., 2017). Previous neuroimaging studies investigating semantic integration in sentence endings with different levels of expectancy (cloze probability) showed increased activity in key areas of the semantic network, including the left anterior inferior frontal gyrus (aIFG), posterior superior temporal sulcus/middle temporal gyrus (pSTS/MTG), and anterior superior temporal sulcus/MTG (aSTS/MTG) for sentence endings with high integration demands, such as unexpected and anomalous endings (Baumgaertner et al., 2002, Hartwigsen et al., 2017).

To elucidate the mechanisms underlying semantic integration, it is essential to briefly explore the field of semantic cognition research. In the current literature, semantic cognition is typically defined as the effective use of acquired knowledge about the world (Badre & Wagner, 2002; Hoffman et al., 2018; Jefferies, 2013; Lambon Ralph et al., 2017). A growing body of evidence suggests that successful semantic cognition is based on two interacting components (Jefferies, 2013; Lambon Ralph et al., 2017): 1) Semantic knowledge representation refers to the ability to store information about the meaning of objects, concepts, and words. 2) Semantic control processes regulate how aspects of our knowledge are retrieved and used in a specific context or during a specific task (Hoffman et al., 2018; Yee & Thompson-Schill, 2016). All semantic tasks require the interaction of knowledge and control processes, although the weighting between both components may vary depending on the task. Moreover, studies investigating semantic cognition in the lesioned (e.g., Jefferies & Lambon Ralph, 2006) or aging brain (e.g., Hoffman, 2018; Hoffman & MacPherson, 2022) suggest that both aspects can be impaired independently and rely on different neural systems. For example, older people generally show well-preserved semantic knowledge, whereas semantic control processes deteriorate with age (Grady, 2012; Hoffman, 2018; Hoffman & MacPherson, 2022; Martin et al., 2022, 2023; Morcom & Johnson, 2015; Wu & Hoffman, 2022). At the neural level, several studies have shown that the anterior temporal lobes function as a store of conceptual representations and are largely responsible for semantic knowledge through interaction with modality-specific association regions (Hoffman et al., 2017; Humphreys et al., 2015; Mion et al., 2010; Patterson et al., 2007; Pobric et al., 2007). The controlled use of semantic knowledge, on the other hand, is primarily associated with the activation of a neural network that includes the inferior frontal gyrus, the posterior middle temporal gyrus (pMTG), and inferior parietal regions including the intraparietal sulcus (Noonan et al., 2013; Rodd et al., 2005; Thompson-Schill et al., 1997).

Since these brain regions align well with the above discussed regions involved in semantic integration under increasing demands, this leads to the assumption that higher semantic integration demands require enhanced control processes within the semantic network. Several studies have investigated network connectivity during the performance of semantic tasks using fMRI (e.g., Davey et al., 2016; Jackson et al., 2016) or MEG (e.g., Kielar et al., 2016). However, it is less clear how exactly brain areas interact during semantic integration. Hartwigsen et al. (2017) explored task-related effective connectivity during semantic integration between preselected areas of the semantic network. This study revealed task-specific interactions between temporal and frontal areas: Relative to sentences with expected endings, sentences with unexpected endings increased the inhibitory influence of the left aSTS/MTG on left pSTS/MTG. In contrast, processing sentences with semantically anomalous endings required increased inhibitory connectivity from left aIFG to left pSTS/MTG. These results are supported by other studies that complemented task-related interactions during semantic processing with resting state connectivity (e.g., Jackson et al., 2016; Mascali et al., 2018). For example, Wawrzyniak et al. (2017) revealed a strong interaction between predefined areas in the left IFG, aMTG, and pMTG, and a left-dominant frontotemporal network of semantic language regions at rest. Increased functional connectivity between IFG and pMTG correlated positively with task performance under high semantic integration demands in the task paradigm. These results emphasize the relevance of interactions between the left IFG and pMTG in semantic control processes.

Previous research has shown that, in addition to a functionally specialized core network, domain-general networks are also crucial for successful language processing (e.g., Fedorenko & Thompson-Schill, 2014). Indeed, tasks with high semantic control demands led to increased activation in areas that partially overlap with the multiple demand network - a network activated in response to increased executive control demands across various cognitive domains (Duncan, 2010; Fedorenko et al., 2013). FMRI studies suggest that semantic control can be divided into semantic-specific and domain-general components that activate adjacent brain regions in the left inferior prefrontal cortex (Badre et al., 2005; Hoffman, 2018; Nagel et al., 2008). Controlled retrieval appears to be semantic-specific, a mechanism that comes into play when automatic activation of semantic knowledge is insufficient, e.g., when a less dominant or unexpected word meaning must be accessed (Badre & Wagner, 2007). This controlled retrieval is associated with increased activation in the anterior ventral part of the left inferior prefrontal cortex (Brodman Area 47 (BA47)) (Badre et al., 2005; Dobbins & Wagner, 2005; Gold et al., 2006; Jackson et al., 2016; Krieger-Redwood et al., 2015). Studies of the structural and functional connectivity of BA47 support this hypothesis, as BA47 has close connections to anterior temporal regions, i.e., the region where semantic knowledge is stored (Jackson et al., 2016; Jung et al., 2017; Von Der Heide et al., 2013). In contrast, semantic selection between competing representations seems to be controlled more by a domain-general executive selection system (Hoffman, 2018). Here, increased brain activation has been found primarily in the posterior part of the left inferior prefrontal cortex (BA 44/45) (Badre et al., 2005; Gold et al., 2006; Thompson-Schill et al., 1997). Therefore, the interaction between domain-specific semantic regions and domain-general networks likely contributes to semantic integration. As the previous study by Hartwigsen et al. has primarily focused on effective connectivity between pre-selected areas of the semantic network during semantic integration, it is unclear how varying semantic integration demands modulate functional connectivity between large-scale cognitive networks at the whole-brain level. The present study aims to address this gap. Specifically, we were interested in whole-brain interactions during the processing of sentences with unexpected and anomalous endings.

Using a previously established auditory semantic integration paradigm (Hartwigsen et al., 2017), we combined univariate task-related functional magnetic resonance imaging (fMRI) analyses with seed-based connectivity and between-network connectivity analyses. This allowed for a comprehensive characterization of semantic integration at the whole-brain level. We expected that semantic integration demands should modulate the task-specific interaction between frontal, temporal, and potentially parietal areas. More specifically, we hypothesized that processing sentences with increased semantic integration demands, that is, sentences with unexpected and anomalous endings, would lead to a stronger interaction between brain regions involved in semantic control in the frontal and temporal cortex. We assumed that while processing unexpected sentence endings requires the suppression of the expected word, semantic integration generally succeeds. In contrast, for anomalous sentence endings we hypothesized that, while lexical word retrieval should be possible, semantic integration would require restructuring attempts. Consequently, we expected to find an even more pronounced interaction with frontal control regions during the processing of anomalous sentence structures compared to unexpected ones. Additionally, we examined the processing of sentences with pseudoword endings, where neither lexical word retrieval nor semantic integration of the pseudoword into the sentence context should be feasible. Here, we also anticipated a significant interaction with control regions, even though this condition likely leads to less pronounced recruitment of the semantic system compared to sentences with real word endings. In addition to the increased interaction between regions of the semantic network, we hypothesized that the interaction with domain-general networks, particularly during the processing of sentences with increased integration demands, is crucial. We expected that the interaction between task-positive networks increases as semantic integration demands rise. Conversely, we expected a stronger interaction with the default-mode network (DMN) during the more automated processing of predictable sentence endings. At the behavioral level, increased semantic integration demands should be reflected in increased response latencies and probably also decreased accuracy. Specifically, we expected an increase in response speed from expected to unexpected, anomalous and pseudoword endings. Furthermore, we assumed that behavioral performance might benefit from a stronger coupling between task-positive networks and a decreased coupling with task-negative networks such as the DMN.

## Methods

### Participants

33 healthy participants were recruited via postings at the University Hospital Halle (Saale) and the database of the Max Planck Institute for Human Cognitive and Brain Sciences in Leipzig. One participant was excluded because of left-handedness. The final sample included 32 participants (mean age: 55.7 years, SD: 13.2, range: 30-75 years, 15 females). For a visualization of the age distribution, see **Supplementary Material Figure S1**. On average, the participants had 16 years of education (SD: 2.4). All participants were native German speakers and, according to the Edinburgh Handedness Inventory (Oldfield, 1971), right-handed (mean LQ: 91.5, SD: 9.0). They had normal hearing, normal or corrected-to-normal vision, and no history of neurological or psychiatric conditions or contraindications to magnetic resonance imaging (MRI). The study was approved by the local ethics committee of the University of Halle (Saale) and conducted in accordance with the Declaration of Helsinki. Participants gave written informed consent prior to the experiment. They received 10 Euro per hour for participation.

### Neuropsychological Assessment

To assess cognitive functioning, all participants performed a comprehensive neuropsychological test battery. They were screened for cognitive impairments using the Mini Mental State Examination (MMSE; Folstein et al., 1975; all ≥ 26/30 points) and for depression with the Beck Depression Inventory (BDI-II; Beck et al., 1996; all ≤ 13 points). Furthermore, participants performed the Digit Span Test (Wechsler & De Lemos, 1981) to assess their working memory capacity. The participants also underwent the subtests Alertness, Go/No-go 1 of 2, and Go/No-go 2 of 5 of the Test of Attentional Performance (TAP; Zimmermann & Fimm, 2002). This battery evaluates different aspects of attention, including vigilance, response inhibition, and sustained attention. Demographic data and neuropsychological test results are shown in the **Supplementary Material (Table S1)**.

### Experimental Design and Stimuli

Our study employed an auditory cloze probability paradigm as previously described (Hartwigsen et al., 2017). This paradigm allows for modulating the demands of semantic integration by varying the predictability of the last word in a sentence across four different experimental conditions. The final word can either be (1) expected (e.g., “The pilot flies the *plane*.”), (2) unexpected (e.g., “The pilot flies the *kite*.”), or (3) semantically anomalous (e.g., “The pilot flies the *book*.”). Additionally, pseudoword endings (e.g., “The pilot flies the *kirst*.”), make successful semantic integration impossible. Participants had to perform a lexical decision task (word or a pseudoword) on the sentence’s final words. Stimuli consisted of 60 sentences for each word condition (expected, unexpected, anomalous words), and 120 sentences for the pseudoword condition (i.e., 60% word endings vs. 40% pseudoword endings, 300 sentences in total). To ensure that the context matched between the four conditions, the same sentence stems (subject + verb phrase) were used, with only the last word (object phrase) being varied accordingly. In addition to the 300 experimental trials, a separate set of 50 practice sentences was used for training outside of the scanner. All sentences were recorded by a female, professional German speaker. For details regarding stimulus creation, please refer to Hartwigsen et al. (2017).

### Experimental Procedure

The data were collected in a single session of three hours for each participant. After the participants provided informed consent, they completed the neuropsychological assessment, followed by a short training of the experimental task outside the scanner. The subsequent MRI measurement lasted about one hour and included the acquisition of a T1-weighted anatomical dataset as well as several fMRI scans (resting state, semantic integration paradigm, language localizer, breath hold task). Here, we focus on the results of the semantic integration task, as the other measures were performed to collect control data in the context of an ongoing patient study. The semantic integration task was implemented in two event-related fMRI runs and had a total duration of 27 minutes. During the experiment, participants looked at a black screen with a white fixation cross in the center. All 300 stimuli were presented auditorily via MR-compatible over-ear headphones. Sentence duration ranged from 1.5 - 2.6 s, with a jittered inter-trial interval of 1.5 - 4.4 s. Both runs contained the same number of stimuli for each condition (30 expected, 30 unexpected, 30 anomalous, 60 pseudowords). The distribution and order of stimuli across both runs were generated for each participant individually as a pseudo-randomized list, ensuring that a maximum of three sentences from the same condition could occur consecutively. Subjects performed a lexical decision task (Is the last word a real word or a pseudoword?) by responding via button press with their left middle (pseudoword) or index finger (word), and reaction times and accuracy data were collected. Note that the left hand was used to avoid button-press related left-hemispheric motor activity. Presentation of the stimuli and triggering of the scanner was conducted through PsychoPy (Peirce et al., 2019). For a visualization of the experimental design see **Figure 1**.

**Figure 1.**
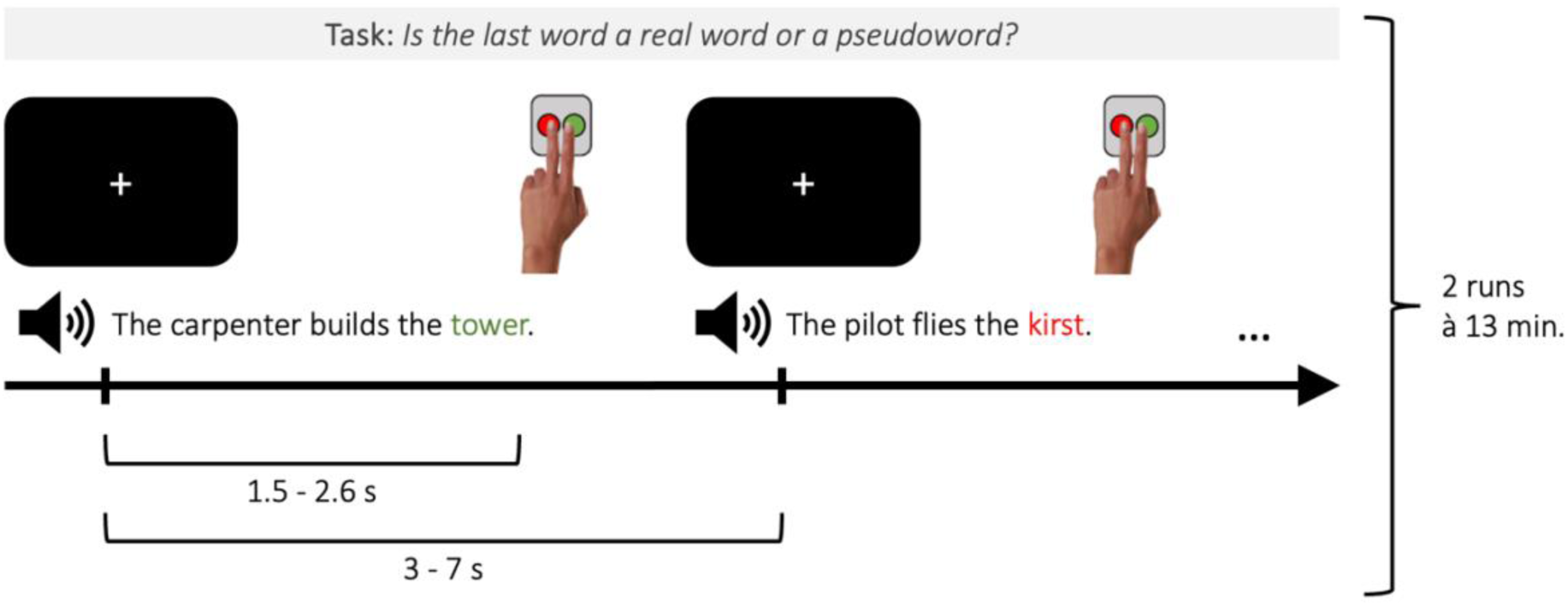
Experimental design of the semantic integration task. The figure illustrates two fMRI trials. Sentences were presented in an event-related fashion with a variable stimulus onset asynchrony between 3 and 7 s. Sentence duration varied between 1.5 and 2.6 s. After the auditory presentation of the sentence, subjects had to indicate via button press whether the final word represented a word or pseudoword (lexical decision). Reaction times (from the last word onset) and error rates were measured.

### Data Acquisition and Preprocessing

Functional imaging was performed on a human whole body 3-Tesla scanner (Magnetom Skyra, Siemens, Erlangen, Germany) equipped with a 32-channel head coil. For the acquisition of fMRI data during the semantic integration task, a gradient echo-planar imaging (EPI) sequence was used (TR/TE = 2.24 s / 0.03 s, flip angle 90°, field of view (FOV): 100 mm, matrix: 76×76 pixel, voxel size: 3×3×3 mm). A total of 360 image stacks consisting of 40 transversal slices each were acquired continuously during each session. Additionally, T1-weighted anatomical images were acquired with an MPRAGE sequence in sagittal orientation (voxel size = 1 x 1 x 1 mm; TR = 2.53 s; TE = 0.00267 s). Preprocessing was performed using fMRIPrep 23.0.0 (Esteban et al., 2019), which is based on Nipype 1.8.7 (Gorgolewski et al., 2011). In summary, preprocessing comprised skull stripping, co-registration, slice timing correction, and calculation of several confounding time-series for each of the two functional runs per participant. Anatomical T1-weighted images were skull-stripped, segmented, and spatially normalized to the MNI152NLin6Asym template. For details on the preprocessing pipeline, see **Supplementary Material (Note S1)** and fMRIPrep’s documentation (https://fmriprep.org/en/23.0.0/workflows.html). Finally, functional images were smoothed with a 6 mm FWHM Gaussian kernel using SPM12 implemented in MATLAB R2022b (9.13; The Mathworks Inc., Natick, MA, USA).

### Behavioral Data Analysis

The statistical analysis of reaction times and error rates was conducted using R 4.2.2 via RStudio (R Core Team, 2021). We utilized the packages lme4 (Bates et al., 2014) and ggeffects (Lüdecke, 2018) for mixed models and ggplot2 (Wickham, 2016) for visualizations. Reaction times were measured from the onset of the last word in a sentence to the button press. Incorrect responses, omissions, and trials in which the button was pressed before the onset of the last word were excluded from the analysis of reaction times. Reaction times were corrected for outliers (based on the individual means by subject and condition +/- 2 SDs). For the analysis of reaction time data (**Equation S1**), a linear mixed-effects model with the log-transformed data was computed. As fixed effects, we included condition, age, and their interaction term. Intercepts for participants and stimuli were defined as random effects. Additionally, we entered education as a covariate of no interest to account for a potential effect on performance in the semantic integration task. P-values were calculated using likelihood ratio tests comparing the full model with the effect in question against the model without the effect in question. The emmeans package (Lenth, 2020) was applied for post hoc comparisons, using Bonferroni-Holm correction. For the analysis of error rates, a generalized linear mixed-effects logistic regression was used to account for the binary nature of the response variable (**Equation S2**). However, since the model including an interaction of condition and age failed to converge, we defined the model without this interaction.

We used deviation (simple) coding for our categorical predictor “condition”. The purpose of simple coding is to create numeric variables that capture deviations from the overall mean of a continuous variable or the baseline level in case of categorical variables, thereby allowing for comparisons in terms of deviations from a central reference point across different levels of a variable. The numeric variables “education” and “age” were mean-centered and scaled to have unit variance.

### Univariate fMRI Analysis

For the statistical analysis of fMRI data, a two-level approach was implemented using SPM12. On the first level, a general linear model (GLM) with a flexible factorial design was created for each participant. Each run was modeled separately in this design. The GLM included regressors for the onsets and durations of the four stimulus conditions, as well as a regressor of no interest for incorrect trials (wrong button presses). Nuisance regressors, consisting of the six motion parameters and individual regressors for strong volume-to-volume movement, as indicated by framewise displacement (FD) values > 0.9, were also defined. Additionally, age was included as a covariate in the model. A high-pass filter with a cut-off of 128 s was applied to the data before model estimation. Statistical parametric maps of the t-statistic were generated by estimating the contrast of each condition against rest and the direct contrasts between the individual conditions. Additionally, the contrasts “sentences with word endings > sentences with pseudoword endings” and “sentences with pseudoword endings > sentences with word endings” were estimated by combining the trials of the three word-conditions.

For the second-level analysis, the contrast images of the pooled parameter estimates were entered into a random effects model. A one-sample t-test was performed for within-group comparisons. Contrasts were thresholded at p < 0.05 and corrected for multiple comparisons using the family-wise error method at the cluster level (FWEc). The SPM anatomy toolbox (version 3.0; Eickhoff et al., 2005) was used for the anatomical localization of activation peaks.

### Functional Connectivity Analysis

#### Generalized Psychophysiological Interaction Analyses

To investigate task-related changes in functional connectivity during semantic integration, we conducted a generalized psychophysiological interaction (gPPI) analysis using the gPPI toolbox for SPM12 (version 13.1; McLaren et al., 2012). Since we were mainly interested in the functional connectivity underlying challenging but successful semantic integration processes, we defined our seed regions for areas that showed greater activation for unexpected and anomalous sentence endings relative to expected sentence endings. To this end, we ran a conjunction analysis: “unexpected > expected ∩ anomalous > expected” (see **Figure S2** in the **Supplementary Material**). Subsequently, seed regions were defined for all global maxima within this conjunction (FWEc, p < 0.05) (see **Table S2**). A spherical ROI with a radius of 10 mm was created around each peak coordinate, using the MarsBaR toolbox (version 0.45; Brett et al., 2002). Within this boundary, we searched for the top 25% active voxels (positive) in each participant and defined them as our subject-specific ROIs. All participants had active voxels within the spherical masks. Next, we conducted the standard two-level approach by performing a whole-brain random-effects group analysis based on the GLM. At the first level, individual participant data were modeled separately using the gPPI toolbox. The first level GLM included: 1) “Psychological” regressors for all four experimental conditions convolved with the canonical hemodynamic response function (HRF), 2) a “physiological” regressor consisting of the deconvolved time series of the first eigenvariate of the BOLD signal from the respective seed ROIs, 3) PPI regressors for each experimental condition, which were calculated by multiplying the deconvolved BOLD signal of the respective seed ROIs with the condition onsets and convolving with the canonical HRF (McLaren et al., 2012), 4) nuisance regressors, consisting of the six motion parameters and individual regressors for strong volume-to-volume movement, as indicated by FD values > 0.9. Resulting participant level contrast images were entered into t-tests on the group level. To test for functional coupling during semantic integration, we compared the connectivity for all our task contrasts by using paired t-tests. For all group-level analyses, a gray matter mask (SPM12 tissue probability map) was applied, restricting statistical tests to voxels with a gray matter probability > 0.3. All activation maps were thresholded at a voxel-wise p < 0.001 and a cluster-wise p < 0.05 FWE-corrected for multiple comparisons. We also tested the gPPI analysis using seed regions defined by the peak coordinates of the individual contrasts unexpected > expected and anomalous > expected (instead of the conjunction of both). However, no significant results were found for this approach.

#### Independent Component Analysis

To further assess which large-scale networks are active during the semantic integration task, we conducted a groupwise spatial independent component analysis (ICA; Calhoun et al., 2009). ICA decomposes fMRI time series into multiple source components in a data-driven manner. While artifactual components can be removed (Griffanti et al., 2014), the remaining independent components correspond to functionally connected neural networks.

The preprocessed, normalized, and smoothed data were analyzed using the Group ICA of fMRI Toolbox (GIFT v4.0c), which initially intensity-normalized the data before implementing the ICA. Data dimensions were reduced with a two-step expectation maximization principal component analysis (PCA) procedure. To this end, PCA was first performed at the run level, with dimensions being reduced from the full time-course-length (360 timepoints) to a participant-specific number of components as determined by the Minimum Description Length (MDL) criterion (Rissanen, 1978). Second, data were concatenated to further reduce dimensionality on the group level, again using the MDL criterion to determine the number of independent components (ICs). The MDL approach allows for an adaptive determination of the optimal number of components based on the data characteristics, providing a more tailored analysis compared to a fixed number of components (Rissanen, 1978; Schwartz, 1978). ICA was implemented using the Infomax algorithm (Calhoun et al., 2002). To ensure the reliability of the decomposition, Icasso was repeated 50 times with a cluster size determined by the MDL-derived number of components (Himberg & Hyvarinen, 2003). In the last step, the back-reconstruction of group-level ICs to the subject level was performed using the GICA3 algorithm in GIFT (Calhoun et al., 2001). For the remaining analyses, the components were scaled to Z-scores within each component. The resulting 36 ICs were visually inspected following the established criteria of Griffanti et al. (2014). After discarding components attributable to artifacts and noise, we identified a total of 11 network components. The spatial extent of these 11 components at the group level was determined using a one-sided t-test on the spatial maps of the participants. Results were corrected for multiple comparisons using a cluster-level threshold at p < 0.05 with the family-wise error (FWE) method.

#### Brain Network Identification

To guide visual inspection in determining network labels for the 11 resulting component maps, we relied on the Jaccard similarity coefficient (*J*; Jaccard, 1912). The Jaccard Similarity Coefficient computes spatial similarity in the form of overlapping voxels between two binary spatial network masks (i.e., a component map A and a template map B), compared to all other voxels in the brain. This spatial similarity measure results in values between 0 and 1, where 0 indicates no similarity, and 1 indicates a complete match. As template images, we chose the 17-network functional connectivity-based parcellation scheme by Yeo et al. (2011) which is based on resting state data and provides a well-established foundation for the functional parcellation of the brain, enhancing the comparability and reproducibility of results.

#### Network Activity

To investigate the activity of networks during various conditions of the semantic integration task, we used the temporal sorting utility in GIFT. Multiple regression analyses were performed between the time courses of the selected components and the design matrix from the GLM at the subject level. Through this analysis step, beta values for each network and each condition were obtained per run, representing the activity of a network for a given task predictor (11 ICs x 4 conditions x 32 subjects x 2 runs).

Initially, we examined the FDR-corrected network activity for each individual task condition compared to the rest condition. We decided to retain only those networks that exhibited significant (de)activation (p < 0.05) for at least one condition versus rest for further analysis. This resulted in a selection of seven networks and ensured that our subsequent analysis of differences in network activity between conditions focused only on networks that seemed to be task relevant.

To determine how the seven networks responded during individual conditions, pairwise comparisons of activity estimates for the respective conditions were conducted. Since this study focused on investigating semantic integration under challenging conditions, we were particularly interested in network activity during unexpected and anomalous compared to expected sentence endings. Additionally, we examined network activity for the two most opposing conditions: pseudoword endings (which cannot be semantically integrated into the overall meaning of the sentence) compared to expected sentence endings (which can be most easily integrated). To this end, the primary beta values for each network were initially averaged across runs. Subsequently, the utility in GIFT, “Stats on Beta Weights,” was used to conduct paired t-tests on the subject level for the chosen conditions (unexpected vs. expected, anomalous vs. expected, pseudoword vs. expected). The results were FDR-corrected, and the significance threshold was set at p < 0.05.

#### Network Interactions

To investigate task-related interactions between the seven networks that showed significant network activity in the task vs. rest comparison, we applied a correlational psychophysiological interaction (cPPI) analysis (Fornito et al., 2012). CPPI computes pairwise partial correlations between ROIs and produces an output in the form of undirected, symmetrical connectivity matrices. To examine network interactions for the tasks reflecting increased semantic integration demands, we performed cPPI for the contrasts “sentences with unexpected vs. expected endings” and “sentences with anomalous vs. expected endings”. Additionally, we were interested in contrasting sentences with semantically non-integrable pseudoword endings and sentences with easily integrable, expected word endings.

During cPPI analyses, the deconvolved time series of each network were multiplied with the task time course from the design matrix of the first-level analysis and convolved with a canonical HRF to create a PPI term. Subsequently, pairwise partial correlations between the PPI terms and two networks were estimated, while controlling for all remaining regressors in the GLM, the noise regressors, and the activity of the ten remaining networks. Task-specific network interactions were assessed by contrasting the above-mentioned conditions of interest. As a result, a symmetrical 7 x 7 connectivity matrix was obtained for each subject and contrast. Finally, the correlation coefficients were Fisher-transformed into z-values and the full matrices, including positive and negative correlation weights, were statistically tested at the group level. Interactions were considered significant at p < 0.05. Additionally, we used the Network-Based Statistic (NBS) Toolbox (Zalesky et al., 2010) to investigate potential differences in the coupling of networks between the task contrasts. However, no significant results were found in this analysis.

#### Behavioral Performance and Network Interactions

For the contrasts “unexpected vs. expected sentence endings,” “anomalous vs. expected sentence endings,” and “pseudoword vs. expected sentence endings,” we investigated the effects of the interaction among networks on participants’ reaction times and error rates by fitting linear mixed-effects models (**Equations S3 and S4**). Models included fixed effects for the corresponding mean-centered network measure, condition, their interaction term, age, education and by-participant random intercepts.

Additionally, we investigated the performance of the participants in the neuropsychological tests with regard to a potential correlation with the connectivity values of the networks. For this purpose, a sum score from all neuropsychological test results was calculated for each participant. Subsequently, a correlation analysis between the sum scores and the connectivity values of the networks for the respective task contrasts was conducted. However, after removing an outlier, no significant correlations were observed here.

## Results

### Semantic integration demands are reflected in behavioral responses

Analyses of reaction times revealed a significant two-way-interaction of condition and age (χ2 = 24.3249, p < 0.001). Post hoc tests showed significant differences in reaction times between all three word conditions (all p < 0.001), with the shortest reaction times observed for sentences with expected endings, followed by sentences with unexpected and anomalous endings. The longest reaction times were measured for sentences with pseudoword endings. However, reaction times for anomalous vs. pseudoword sentence endings did not differ significantly (**Figure 2, left**). For error rates, we found main effects of condition (χ2 = 71.793, p < 0.001) and age (χ2 = 8.018, p < 0.001). Post hoc tests revealed significant differences in error rates between all conditions (all p < 0.001), except for the contrast of unexpected vs. expected sentence endings (p = 1.0). The highest error rate was found for sentences with anomalous endings, followed by sentences with pseudoword endings and unexpected endings. The lowest error rate was observed for sentences with expected sentence endings (see **Figure 2, right**). Complete model outputs are reported in the **Supplementary Material (Table S3)**. Furthermore, both reaction times and error rates, particularly for conditions with increased semantic integration demands, showed an increase with advancing age (see **Figure S3**).

**Figure 2.**
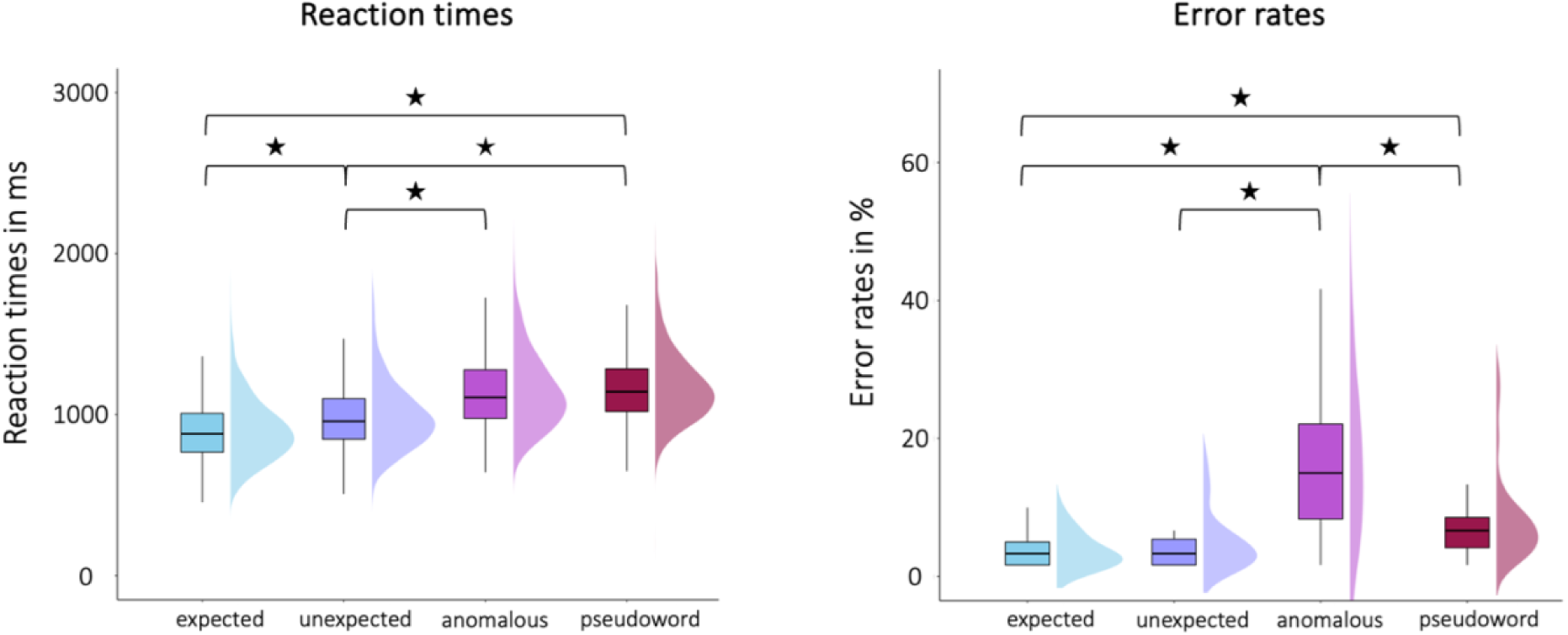
Behavioral results. Differences in reaction times (RTs, left panel) and error rates (right panel) between task conditions. *p < 0.05, Bonferroni-Holm corrected.

### Semantic integration demands increase task-related activity in left fronto-temporal areas

Our goal was to investigate the neural correlates underlying semantic integration under varying conditions. For all examined contrasts, we observed the expected brain activation in a predominantly left-lateralized fronto-temporo-parietal language network (**Figure 3 and Table S4**).

**Figure 3.**
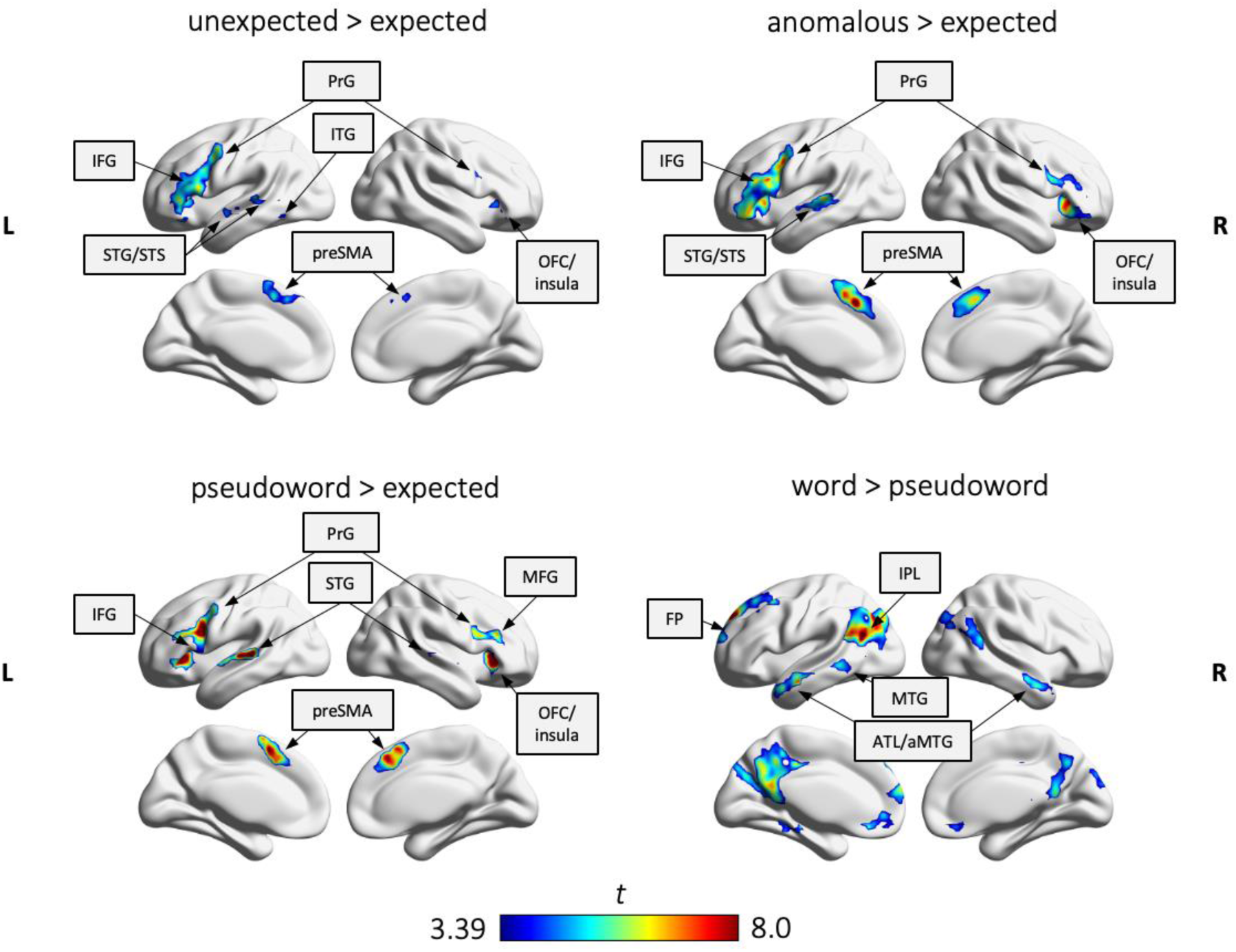
FMRI results on the group level from the univariate analysis for our semantic contrasts of interest. Results are FWE-corrected at p < 0.05 at the cluster level. Abbreviations: ATL: Anterior temporal lobe; OFC: Orbitofrontal cortex; FP: Frontal pole; IFG: Inferior frontal gyrus; IPL: Inferior parietal lobe; ITG: Inferior temporal gyrus; (a)MTG: (Anterior) middle temporal gyrus; PreSMA: Presupplementary motor area; PrG: Precentral gyrus; STG: Superior temporal gyrus; STS: Superior temporal sulcus.

Increased semantic integration demands were generally reflected by a frontal activation pattern. Specifically, the contrasts of unexpected, anomalous and pseudoword sentence endings vs. expected sentence endings all showed activation in the inferior frontal gyrus (IFG), the orbitofrontal cortex (OFC), and the presupplementary motor area (preSMA). For the “unexpected sentence endings > expected sentence endings” contrast, additional activation was found in the right precentral gyrus (PrG), the right cerebellum (VI), and the left inferior temporal gyrus (ITG). A similar, although even more pronounced activation pattern emerged for the “anomalous sentence endings > expected sentence endings” contrast. Here we found additional brain activation in the right cerebellum (VI), the left superior temporal gyrus (STG) and the right cerebellum Crus II. Finally, for sentences with pseudoword endings compared to expected endings we observed additional activation in the middle frontal gyrus (MFG).

In contrast, the processing of sentences with expected endings was associated with predominantly temporo-parietal brain activity. We also examined blood-oxygen-level-dependent (BOLD) signal differences during the processing of sentences with word endings (encompassing the conditions expected, unexpected, and anomalous) > sentences with pseudoword endings. We found the largest differences in brain activation in the left inferior parietal lobe (IPL), the left frontal pole (FP), the left middle temporal gyrus (MTG), the left parahippocampal gyrus, and the right MTG. Additional contrasts between the individual task conditions are included in the **Supplementary Material (Figure S4)**. No significant effect of age was found in our univariate analysis.

### Functional Connectivity Results

#### Semantic integration demands increase task-related interactions between frontal and subcortical areas

We ran gPPI analyses for seeds derived from the conjunction of the tasks with increased semantic integration demand, i.e., unexpected and anomalous sentence endings. Results revealed significant task-specific interactions for the seeds in the left IFG and preSMA. For unexpected > expected words, we found increased functional connectivity between left IFG and the thalamus (**Figure 4, left**). For the contrast expected > anomalous words, left preSMA showed increased coupling with the right cerebellum (**Figure 4, right**). We also performed gPPI analysis for regions with increased activation for individual contrasts of unexpected and anomalous conditions relative to expected words. Results did not indicate significant changes in functional connectivity for these seeds.

**Figure 4.**
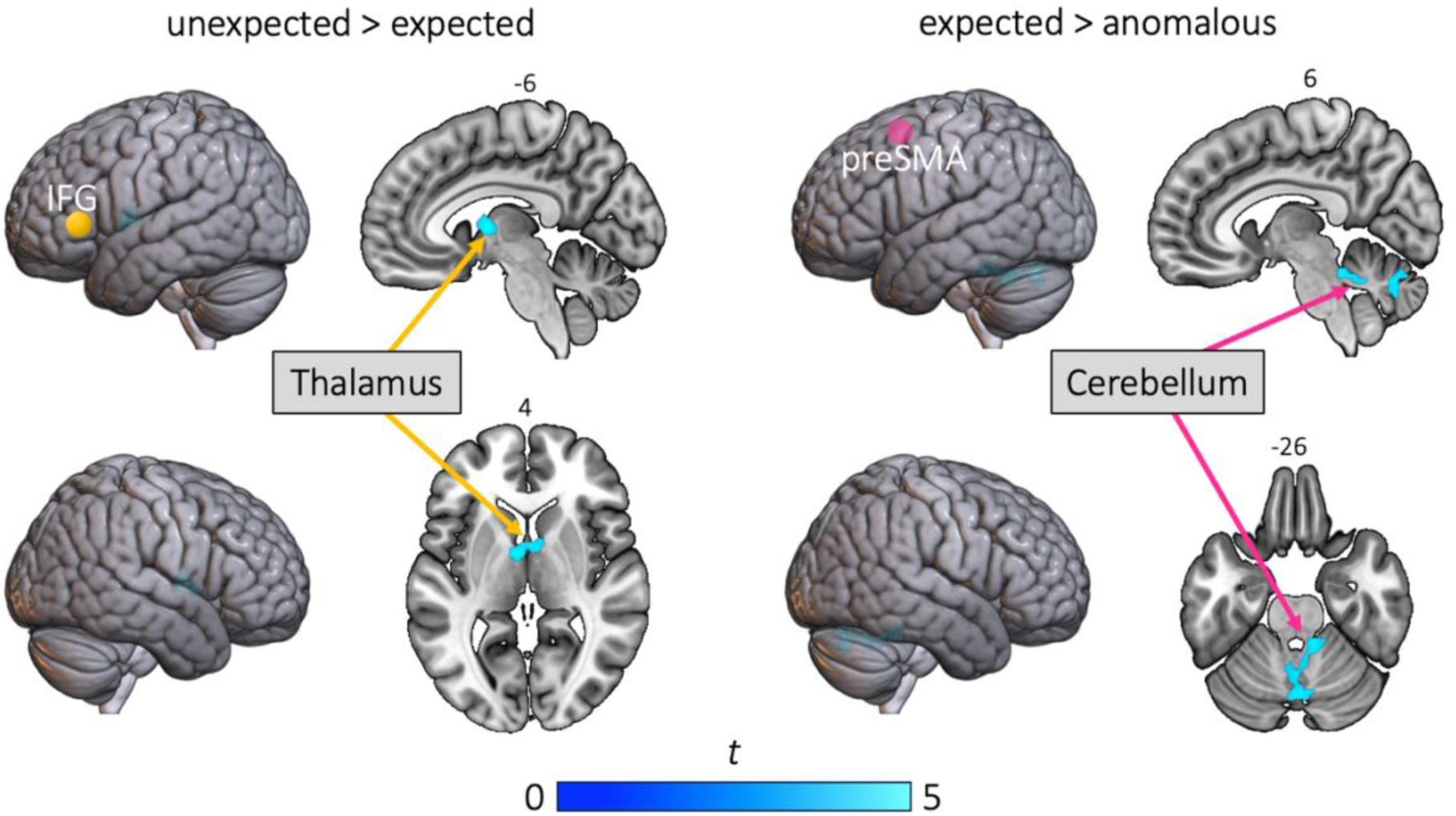
Task-dependent changes in functional connectivity for seeds in the left inferior frontal gyrus (IFG) and the contrast unexpected sentence endings > expected sentence endings (left) as well as the presupplementary motor area (preSMA) and the contrast expected sentence endings > anomalous sentence endings (right). All results are FWE-corrected at p < 0.05 at cluster level.

#### Semantic integration is characterized by distributed large-scale networks

We defined functional networks for the semantic integration task by performing spatial ICA at the group level. After filtering out noise components from a total of 36, we identified 11 network components encompassing ten cortical components and one component in the cerebellum.

To assign the 11 ICs to cognitive networks, we calculated the Jaccard Similarity coefficient between our binarized and thresholded ICs and template masks of neural networks from Yeo et al. (2011). With the exception of IC16 and IC33, all ICs showed a similarity coefficient above the threshold recommended by Jackson et al. (2019) (J = 0.15) for at least one network template. The Jaccard indices for individual ICs and network templates are listed in **Table S5**. IC16 exhibited predominantly cerebellar connectivity and was thus not represented in the cortical templates of Yeo et al. (2011). Consequently, we labeled IC16 as the cerebellum. For all other independent components, we chose the label according to the highest Jaccard similarity coefficient. We found two control networks: ContA (IC25, J = 0.241) and ContB (IC17, J = 0.159), two default mode networks: DefaultA (IC11, J = 0.225) and DefaultB (IC35 = 0.273), as well as two somatomotor networks: SomMotA (IC12, J = 0.274) and SomMotB (IC04, J = 0.288). IC13 showed the highest similarity to the salience/ventral attention network A (SalVentAttnA, J = 0.240). IC18 showed the highest Jaccard similarity coefficient for the central visiual network (VisCent, J = 0.197). Two networks showed the highest agreement with the peripheral visiual network (IC21, J = 0.286; IC33, J = 0.122). **Figure 5** shows the thresholded maps (FWE-corrected at the cluster level with p < 0.05) with their original component numbers. Please note that the figure only includes the selected non-noise components that showed significant (de)activation for at least one task condition, since we decided to focus on those components in our further analysis. Details will be explained in the next paragraph.

**Figure 5.**
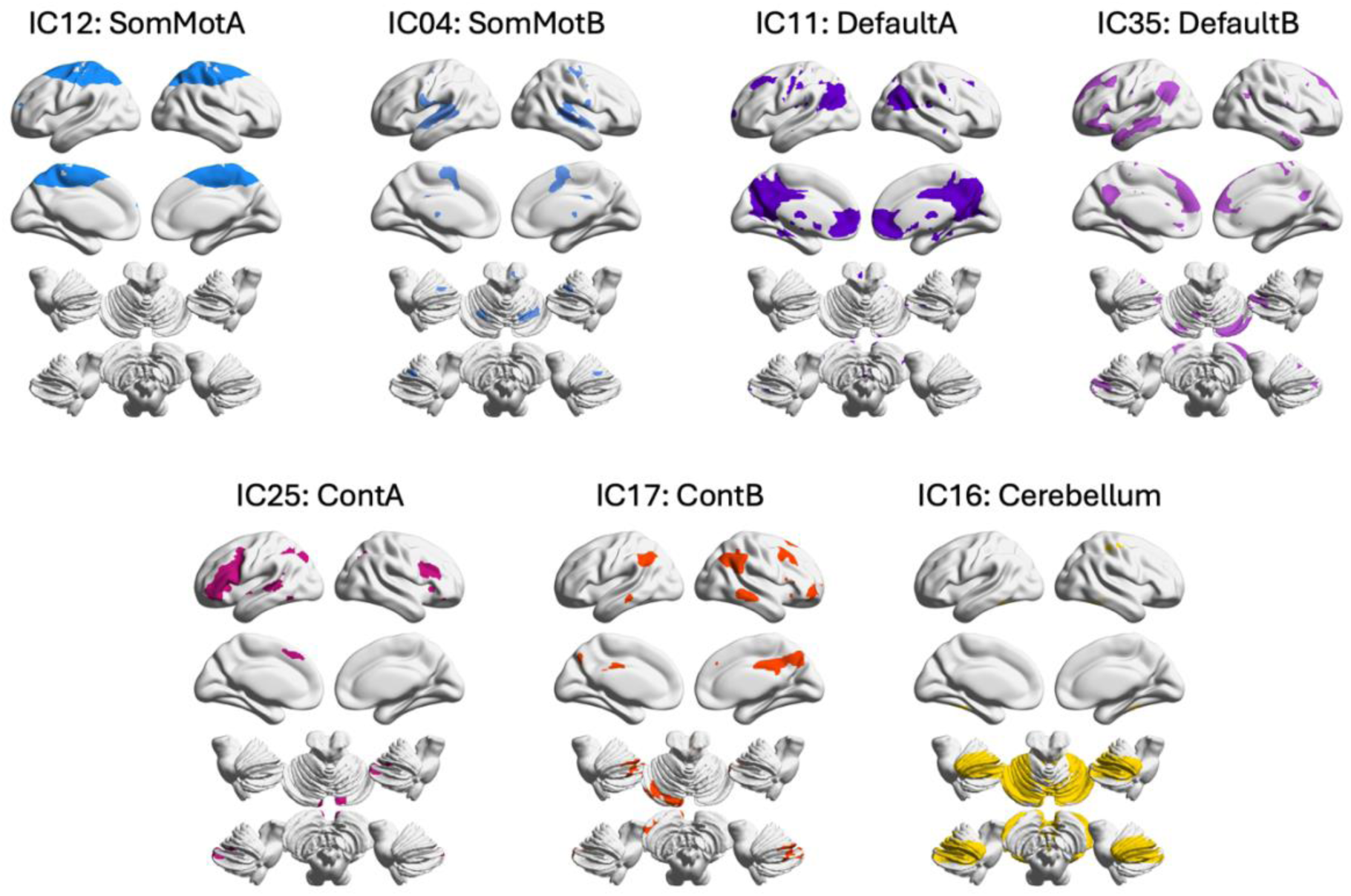
Spatial ICA-derived networks during the semantic integration task. Networks are shown as binary masks, created from the T-scores from 1-sided T-tests (FWE-corrected p < 0.05 at cluster level). Network labels were chosen according to the spatial similarity analysis. Abbreviations: ContA/B: Control network A/B; DefaultA/B: Default mode network A/B; SomMotA/B: Somatomotor network A/B.

#### Semantic integration demands differentially modulate network activity

To focus subsequent analyses on networks relevant to the semantic integration task, we examined network activity for each task condition compared to rest. In total, seven of the 11 networks showed significant (de)activation in at least one task condition (see **Figure 6**, top panel). A detailed summary of the beta and p-values of all 11 networks can be found in **Table S6**. Both somatomotor networks (SomMotA and SomMotB) showed significant positive activation for all four task conditions. The two control networks revealed a more complex picture: While ContA showed significant positive network activity for the three conditions with increased integration demands (unexpected, anomalous and pseudoword), ContB was significantly deactivated for the sentences with unexpected word endings. The two default mode networks showed overall more deactivation, which was significant for all four conditions in the DefaultA network, but only for the pseudoword condition in the DefaultB network. Finally, the cerebellum showed positive network activity that reached significance for the anomalous condition. All other networks (IC13, IC18, IC21, IC33) showed no significant (de)activation and were thus excluded from the following analysis.

**Figure 6.**
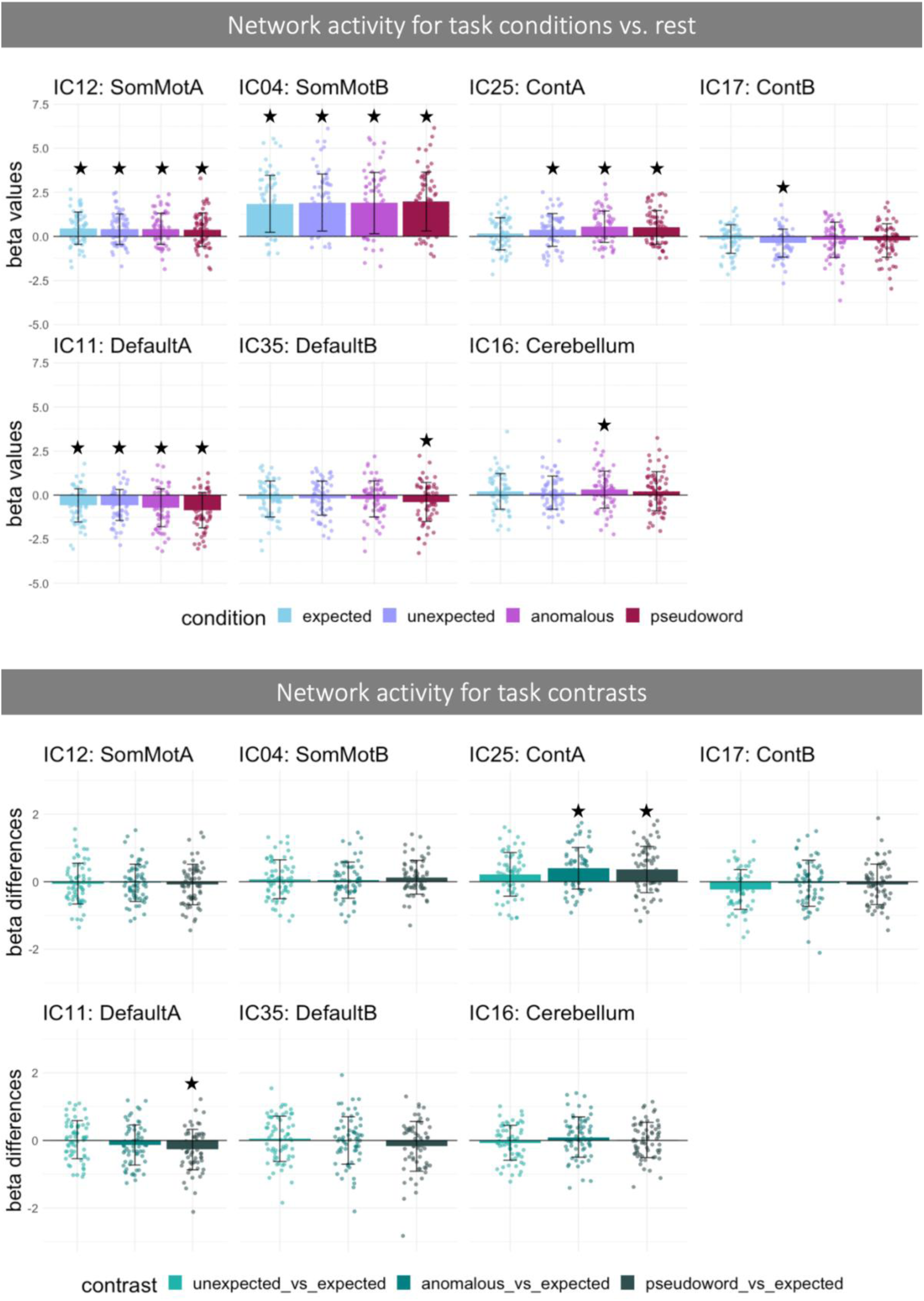
Network activity during semantic integration. Top Panel: Network activity (beta weights ± SD) for the conditions expected, unexpected, anomalous and pseudoword compared to rest. Bottom Panel: Network activity calculated from differences between mean beta weights (± SD) for the contrasts of unexpected, anomalous and pseudoword vs. expected trials. Asterisks indicate significance for a given network’s contrast t-test (FDR-corrected, p < 0.05). Abbreviations: ContA/B: Control network A/B; DefaultA/B: Default mode network A/B; SomMotA/B: Somatomotor network A/B.

Next, we examined the activity of the remaining seven networks during the semantic integration task for three contrasts reflecting increasing integration demands (“unexpected vs. expected sentence endings,” “anomalous vs. expected sentence endings,” and “pseudoword vs. expected sentence endings”). The differences in beta values between conditions are depicted in **Figure 6** (bottom panel) and described in **Table S6** for each network. No significant (de)activation was found for the “unexpected vs. expected sentence endings” contrast. However, a significant positive activation of ContA was detected for the more challenging task contrasts “anomalous vs. expected sentence endings” and “pseudoword vs. expected sentence endings”. Furthermore, DefaultA showed significant deactivation for “pseudoword vs. expected sentence endings”.

#### Semantic integration demands mainly increase interactions between task-positive networks

To investigate how the seven ICA-derived networks interact with each other during different conditions of the semantic integration task, a cPPI analysis was conducted, resulting in a 7 x 7 correlation matrix for each task contrast (**Figure S5 B**). Significant network interactions for our contrasts of interest (“unexpected vs. expected sentence endings,” “anomalous vs. expected sentence endings,” and “pseudoword vs. expected sentence endings”) are shown in **Figure 7** (left column). Additional results are visualized in **Figure S5 A**.

**Figure 7.**
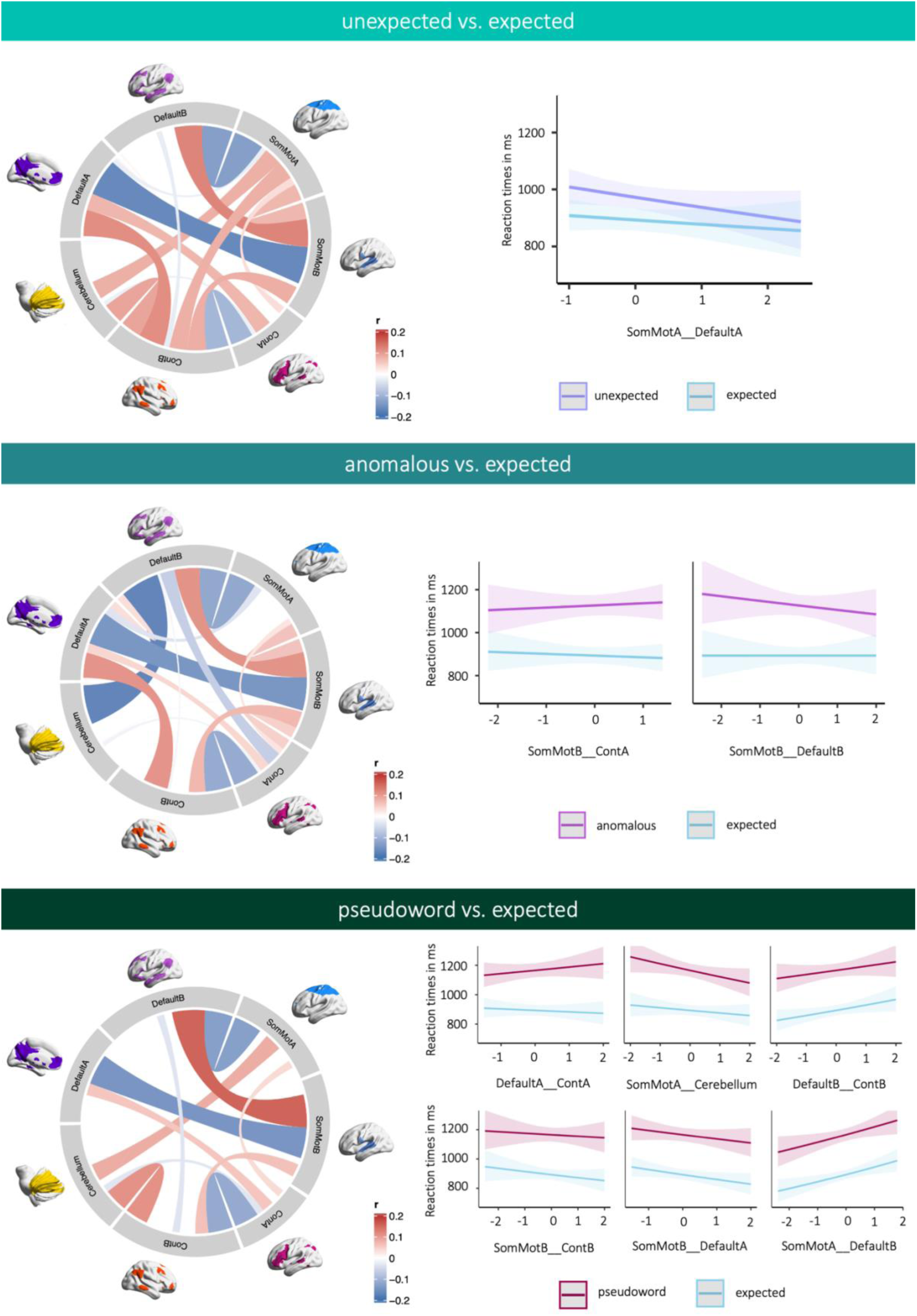
Functional coupling between task-relevant networks and their behavioral relevance. Left panels: Chord diagrams show significant results of functional coupling between ICA-derived networks. Connectivity values are partial correlations. The color intensity and width of a connection indicate its correlational strength. Higher correlation values (r) indicate positive coupling, and negative values indicate decoupling between networks. Right panels: Correlations between network interactions and task efficiency. Plots show significant 2-way interactions between task conditions and the respective network pair for response time data. Connectivity values were mean-centered for interaction analyses. Results were FDR-corrected for multiple comparisons at p = 0.05. Abbreviations: ContA/B: Control network A/B; DefaultA/B: Default mode network A/B; SomMotA/B: Somatomotor network A/B.

Networks typically classified as task-positive (ContA and ContB) as well as somatomotor networks predominantly exhibited increased positive interactions with one another across all three task contrasts. Conversely, these task-positive networks displayed increased negative interactions with default mode networks. Exceptions were the positive interactions between DefaultB and SomMotB, DefaultA and ContA, and DefaultA and ContB, the last pair only for the two semantic contrasts (unexpected vs. expected and anomalous vs. expected), but not for the contrast between pseudowords and expected sentence endings. Furthermore, we found a negative interaction between the two control networks ContA and ContB for all three task contrasts. The cerebellum showed a predominantly positive network modulation with SomMotA and ContB, and a weak negative interaction with ContA. Interestingly, however, this cerebellar interaction pattern differed for the anomalous vs. expected contrast. In this case, an increased negative interaction with DefaultB was predominant, a pattern not observed in the other two contrasts. Furthermore, the results show that for the two semantic contrasts (unexpected vs. expected and anomalous vs. expected), more network interactions reached statistical significance (n = 18) than for the contrast between pseudoword and expected endings (n = 11).

#### Higher integration demands result in an increased number of behaviorally relevant network interactions

Finally, we examined the behavioral relevance of these significant network interactions for the task contrasts “unexpected vs. expected sentence endings,” “anomalous vs. expected sentence endings,” and “pseudoword vs. expected sentence endings” (see **Figure 7, right column**; **Table S7 – S9** for details). We did not detect any significant effect of network coupling on error rates. However, results revealed interactions between reaction times in the respective semantic conditions and network interactions. Remarkably, we found more behaviorally relevant network interactions for higher integration demands. For the “unexpected vs. expected sentence endings” contrast, we found a significant interaction between SomMotA and DefaultA. Increased coupling between these networks was associated with faster reaction times, especially for unexpected trials. For sentences with anomalous endings vs. expected endings, significant network interactions were observed between SomMotB and ContA as well as SomMotB and DefaultB. While a stronger interaction between SomMotB and ContA was reflected in slower reaction times for anomalous trials, coupling led to faster reaction times in the expected condition. The interaction between SomMotB and DefaultB instead promoted faster reaction times for sentences with anomalous endings. Regarding reaction times for sentences with pseudoword endings compared to expected endings, we identified significant network interactions between DefaultA and ContA, SomMotA and Cerebellum, DefaultB and ContB, SomMotB and ContB, SomMotB and DefaultA as well as SomMotA and DefaultB. A stronger coupling between SomMotA and Cerebellum, SomMotB and ContB, and SomMotB and DefaultA resulted in faster reaction times in both conditions. The opposite pattern was found for DefaultB and ContB, as well as SomMotA and DefaultB, since a stronger coupling led to slower reaction times in both task conditions. Finally, a stronger interaction between DefaultA and ContA was associated with slower reaction times in the pseudoword condition and faster reaction times in the expected condition.

## Discussion

The within- and between-network dynamics underlying context-dependent, semantic integration are still poorly understood. Here, we addressed this issue by combining an auditory semantic integration paradigm with univariate task-related fMRI analyses, seed-based within-network analyses, and ICA-based between-network connectivity analyses. On the behavioral level, we reproduced the typical semantic integration effects, characterized by increased reaction times and reduced accuracy for sentences with higher integration demands (Baumgaertner et al., 2002; Hartwigsen et al., 2017). These changes were underpinned by increased task-related activity in brain areas of the semantic control network. Likewise, seed-based functional connectivity analysis revealed a significant interaction between frontal regions involved in cognitive control and subcortical or cerebellar areas. Finally, we identified seven neural networks active during semantic integration. While increasing integration demands promoted activation of task-positive networks, the default-mode networks showed more deactivation as semantic integration demands increased. As a main novel finding, between-network analyses revealed a dynamic interplay between task-positive networks associated with control and somatomotor functions and the default mode network, supporting the processing of sentences with increased semantic integration demands. Increased integration demands lead to a higher number of behaviorally relevant network interactions, suggesting that between-network coupling becomes more relevant for successful task performance as integration demands rise.

The gradual increase in response times with higher semantic integration demands in our data is in agreement with previous studies (Baumgaertner et al., 2002; Hartwigsen et al., 2017; Wawrzyniak et al., 2017) and likely reflects the suppression of the expected word as well as restructuring attempts. Likewise, a similar pattern for accuracy, but with highest error rates for anomalous trials, is supported by the same previous studies. However, unlike the previous studies that selectively included young participants, we did not find a significant difference in reaction times for sentences with anomalous compared to pseudoword endings. This could be explained by the fact that, in our study, older age was associated with longer reaction times and higher error rates, especially for sentences with anomalous endings.

As expected, univariate results revealed a left-dominant, frontal activation pattern, including IFG, OFC, and preSMA, which increased with higher semantic integration demands (cf. Hartwigsen et al., 2017; Wawrzyniak et al., 2017). These regions are associated with cognitive control functions. While the IFG is linked to domain-specific semantic integration (e.g., Hartwigsen et al., 2017; Zhu et al., 2013) and semantic control (Jefferies, 2013), the preSMA and OFC appear to be involved in domain-general control processes (Brockett & Roesch, 2021; Fedorenko et al., 2013; Jackson, 2021). The gradual increase in activation in these regions during the processing of sentences with unexpected, anomalous, and pseudoword sentence endings suggests that the higher the semantic integration demands, the more control is required to inhibit the expected word. In addition to frontal regions, temporal regions were also involved in processing sentences with increased integration demands. Specifically, we found significant activation in the posterior STG/STS. This finding was expected, as the pMTG/STS region has been implicated in the storage of lexico-semantic information (Lau et al., 2008; Price et al., 1997) as well as lexical retrieval and selection under increased executive demands (Whitney et al., 2011, 2012). For sentences with expected endings, as well as for the contrast between word and pseudoword endings, we observed a predominantly temporo-parietal activation pattern, especially in the left IPL/AG. This is not surprising given that this region is a multimodal convergence zone, linking various semantic features of concepts and information from different modalities (Seghier, 2013). In addition, previous studies confirm the critical role of this region in the processing of more predictable sentences (e.g., Humphries et al., 2007).

The gPPI analysis revealed significant task-specific interactions for the left IFG and the preSMA. Specifically, the IFG showed increased interactions with the thalamus for unexpected > expected endings. The thalamus acts as a hub region for many cognitive functions (e.g., Hwang et al., 2017) including language processing and is responsible for the transfer of lexical information to cortical regions (Fritsch et al., 2022; Nadeau & Crosson, 1997). Accordingly, patients with thalamic lesions often show aphasia with mainly lexical-semantic deficits (e.g., Fritsch et al., 2022; Radanovic & Almeida, 2021). Interestingly, the thalamus seems to be involved especially when automated mechanisms such as semantic priming are not effective (Friederici, 2006; Ketteler et al., 2008) or during difficult language tasks (Alain et al., 2005; Assaf et al., 2006). The lexical-semantic nature of our task, and the fact that we found a significant interaction between the IFG and the thalamus only for sentences with increased integration demands fits well into the bigger picture of current research. Furthermore, the preSMA showed increased interactions with the right cerebellum (Crus I-IV and Crus VI) for expected > anomalous sentence endings. An increasing number of studies suggest that the cerebellum is involved in language processing and often exhibits a crossed activation pattern with the left-dominant cortical language network. Areas Crus I-IV and Crus VI have been associated with semantic and phonological processing (Turker et al., 2023). Since both functions are crucial for the performance of our auditory lexical decision task, the involvement of the cerebellum is not surprising here. Consequently, the gPPI results emphasize the importance of subcortical and cerebellar regions in semantic integration, especially under increased demands.

Using spatial ICA, we characterized 11 higher-order large-scale functional networks active during semantic integration. Cortical networks included two control networks: A left hemispheric network (ContA) as well as a right hemispheric network (ContB). Additionally, we identified the “classical” default mode network (DefaultA) and a subnetwork of the default mode network (DefaultB) which is thought to facilitate access to semantic knowledge (Smallwood et al., 2021). We also found two somatomotor networks: SomMotA likely reflects primarily motor aspects, such as button presses (Feis et al., 2015), and SomMotB, which - consistent with the auditory nature of our task - showed pronounced activation in the auditory cortex. Another network was primarily characterized by cerebellar connectivity. Furthermore, we identified a cortical network that best matched the salience/ventral attention network A (SalVentAttnA), a central visiual network (VisCent), and two networks that best aligned with the peripheral visual network (VisPeri). Overall, these results confirm our hypothesis that both task-positive and task-negative networks are involved in semantic integration. The fact that we found both default mode and control networks is consistent with the observation that networks which are typically anticorrelated during resting state become functionally integrated during semantic tasks (Krieger-Redwood et al., 2016; Martin et al., 2022, 2023).

By examining the response of the networks to increasing semantic integration demands, we identified distinct patterns of activation and deactivation across task conditions. Since we were primarily interested in networks that were relevant for successful performance of the semantic integration task, we excluded all networks that did not show significant (de)activation for at least one task condition compared to rest from further analysis. Consequently, we explored task-related activity and between-network interactions for seven task-relevant networks (Cerebellum, ContA and ContB, DefaultA and DefaultB, as well as SomMotA and SomMotB). Our main finding was that increased semantic integration demands were associated with task-positive activation of the left hemispheric control network (ContA) and both somatomotor networks but led to deactivation of the default mode networks. We found the strongest positive activation across all conditions for SomMotB. This result was expected as the network is mainly characterized by connectivity in the auditory cortex and thus likely reflects the auditory nature of the task. The fact that sentences with pseudoword endings elicited slightly more activity in this network than the other three conditions supports this hypothesis and could be explained by increased listening effort for pseudowords. Accordingly, Kotz (2002) showed that lexical status modulates the hemodynamic response in cerebral regions responsible for auditory word recognition. SomMotA also showed significant network activity for all four conditions, likely reflecting motor response (e.g., Feis et al., 2015) caused by the button presses. The slightly lower activity for sentences with pseudoword endings could be due to the imbalance between word and pseudoword trials. Both default mode networks showed stronger deactivation with higher semantic integration demands, although for DefaultB, this effect only reached significance for sentences with pseudoword endings. This result was expected, as the default mode network is known to play a role in self-referential information processing and monitoring the internal mental landscape (Greicius et al., 2003; Qin & Northoff, 2011). These processes are particularly suppressed during cognitive tasks involving external stimuli with increased demands (e.g., Menon & D’Esposito, 2022), such as sentences with unexpected, anomalous or pseudoword endings. Furthermore, evidence suggests that the default mode network activates during more “automatic” retrieval and conceptual combination tasks (Davey et al., 2016; Lanzoni et al., 2020; Price et al., 2016), such as when processing predictable sentences. The observation that DefaultB showed significant deactivation exclusively for sentences with pseudowords (which do not possess a semantic-lexical entry) is consistent with the assumption that this network contributes to semantic processing (Smallwood et al., 2021). The significant network activity of the left-dominant ContA for unexpected, anomalous, and pseudoword sentence endings, but not for expected sentence endings, likely reflects an increased level of cognitive control under increasing integration demands. ContA, together with ContB, constitutes the frontoparietal control network (FPCN), a network that acts as a flexible hub for cognitive control (Dosenbach et al., 2007; Marek & Dosenbach, 2018; Spreng et al., 2010). It is associated with phasic control and attention (for a review, see Menon & D’Esposito, 2022) as well as error-related activity (Dosenbach et al., 2007). These skills are relevant to semantic integration under increased demands. The fact that we found activity mainly in the left-dominant ContA was expected given the semantic language task and may indicate semantic-specific control mechanisms. In contrast, the right-dominant ContB showed an opposite trend with its significant negative activation for unexpected sentence endings. Although this condition is associated with increased integration demands, it still allows for successful semantic integration (unlike the anomalous or pseudoword condition). Therefore, a possible interpretation could be that this condition relies less on right-dominant, domain-general control compared to sentences with anomalous or pseudoword endings. Additionally, the cerebellum showed significant positive network activity for sentences with anomalous endings. This finding is consistent with our gPPI results, where we found an interaction between the control region preSMA and the cerebellum for the contrast between expected and anomalous sentence endings. Thus, the cerebellum may play a supporting role in semantic processing under increased control demands.

Finally, for a direct comparison between network activity under increased integration demands and more automated semantic integration, we also examined network activity for the contrasts “unexpected vs. expected sentence endings”, “anomalous vs. expected sentence endings”, and “pseudoword vs. expected sentence endings”. Positive network activity survived for ContA when processing sentences with anomalous and pseudoword endings compared to expected endings, highlighting the important role of the left-dominant control network in semantic integration under increased demands. DefaultA showed the only significant negative activity for sentences with pseudoword endings compared to expected endings. This result is consistent with our hypothesis that activity in the default mode network decreases with increasing integration demands.

Finally, cPPI analysis unraveled the dynamic interaction between cognitive networks under varying semantic integration demands. Overall, the results reveal distinct interaction patterns among task-positive networks and the default mode networks across varying semantic integration demands. Notably, task-positive control and somatomotor networks exhibited increased positive interactions across all task contrasts, whereas interactions with default mode networks were predominantly negative. Based on the assumption that the FPCN is crucial for domain-general cognitive control and facilitates the realization of new task states through interactions with other relevant networks (Cole et al., 2013; Marek & Dosenbach, 2018; Power et al., 2011), it is not surprising that we observed strong interactions between both control networks (ContA and ContB) and somatomotor as well as default mode networks, independent of task contrast. These findings are consistent with previous research highlighting the role of task-positive networks in cognitive control and attention during increased task demands, while the default mode network is typically anticorrelated (Fox et al., 2005; Power et al., 2011). More specifically, previous studies reported an anticorrelation between the default mode network and control regions in tasks with low semantic predictability (e.g., Lanzoni et al., 2020) and there is evidence that the default mode network may be less involved in harder relative to easy semantic tasks (Humphreys et al., 2015, 2019; Humphreys & Lambon Ralph, 2015). On the other hand, we also found strong positive interactions between DefaultA and the left hemispheric ContA as well as between DefaultB and SomMotB for all three task contrasts, suggesting that the default mode network might contribute to semantic integration to some extent. In fact, there is evidence that the default mode network activates during a wide variety of cognitive tasks, including episodic, linguistic, social and emotional processes and supports the integration of external and internal information (Smallwood et al., 2021; Spreng et al., 2010). In particular, DefaultB shows a large overlap with the semantic network and might therefore contribute to performing our task, as mentioned above. A functional explanation for the default mode network interacting with other task-positive networks during semantic integration under increased demands could be that it facilitates access to semantic knowledge (Smallwood et al., 2021) and thus makes the detection of unexpected stimuli (Corbetta & Shulman, 2002) more effective. Interestingly, we found a negative interaction between the two control networks (ContA and ContB) across all contrasts. This may indicate a division of labor or specialization within the control networks, where different components are engaged depending on the specific cognitive demands of the task (Dosenbach et al., 2007). A comparison of the network couplings for the individual task contrasts revealed slight, although non-significant, differences in between-network interactions. First, we found the most interactions between task-positive networks for sentences with unexpected vs. expected endings, likely supporting successful semantic integration under increased demands for cognitive control and attention. The “anomalous vs. expected sentence endings” contrast showed less pronounced positive network interactions but instead a notable negative interaction between the cerebellum and DefaultB. Since both the cerebellum and DefaultB are known to contribute to semantic processing (Smallwood et al., 2021; Turker et al., 2023), we interpret this negative interaction as a response to the semantic violations caused by the anomalous sentence ending. Finally, the “pseudoword vs. expected sentence endings” contrast, where semantic integration fails due to the missing lexical entry, exhibited the least statistically significant network interactions. This suggests that the absence of meaningful semantic content in pseudowords leads to reduced engagement of both task-positive and default mode networks.

We also examined the behavioral relevance of network interactions during the processing of sentences with varying semantic integration demands, revealing associations between reaction times and network coupling for each task contrast. Our main finding was that increased integration demands led to a higher number of behaviorally relevant network interactions. This suggests that the coupling becomes more relevant for successful task performance as integration demands increase. In general, stronger network interactions were associated with faster reaction times, especially for sentence endings with increased integration demands. However, for the “anomalous vs. expected endings” contrast, stronger coupling between SomMotB and ContA was associated with slower reaction times in the anomalous condition. A possible interpretation would be that subjects with increased listening effort tend to have a stronger interaction between auditory cortex and semantic control regions, which is reflected in a slower button response. In contrast, for sentences with pseudoword vs. expected endings, stronger coupling between default mode and task-positive networks led to slower reaction times. Given that this is the most challenging condition, this result is expected and fits with the fact that the default mode network is often deactivated during the performance of cognitively difficult tasks (Fox et al., 2005; Power et al., 2011). In summary, our findings highlight especially the role of task-positive control and somatomotor networks in facilitating cognitive processing under increased semantic integration demands.

Our study has several limitations that should be addressed in future work. First, the relatively small sample size in combination with a wide age range makes it difficult to draw generalizable conclusions. Our sample was recruited as an age-matched control group for patients with brain tumors in an ongoing study. In the present study, we were able to show a significant age effect in the behavioral data but not in the univariate fMRI and functional connectivity analyses. Thus, our study provides first insights into semantic integration in humans along a continuous age spectrum. However, more comprehensive findings will require larger cohorts. Moreover, investigating functional connectivity during semantic integration in a larger sample of older adults could be insightful, as there is evidence that cognitive decline is associated with increased between-network connectivity (Chan et al., 2014; Martin et al., 2022, 2023; Setton et al., 2022). Another limitation is that the fMRI methods used here are associated with poor signal in the anterior temporal lobe (ATL). This is critical because the ATL is thought to play an important role in semantic cognition research as a storage area for knowledge representations. Although our univariate results for the contrast between sentences with word endings and pseudoword endings showed significant activation in this area, it is possible that anterior and inferior parts are still missing. A multi-echo/multiband fMRI setup would be useful to investigate the role of this brain region in semantic integration under increased demands, especially with regard to its functional connectivity with the rest of the brain. Nevertheless, our findings pave the way for further research on semantic integration in older and clinical populations.

## Conclusion

The present study explored the dynamic within- and between-network interactions during semantic language processing in healthy adults across a continuous age range. We focused on the role of network interactions in semantic integration under increased demands. Behavioral and univariate data revealed typical semantic integration effects, with increased brain activity mainly in regions associated with semantic control for sentences with higher integration demands. Higher semantic control demands were also reflected by significant interactions between control-related frontal regions and subcortical or cerebellar areas. Between-network interactions further highlighted the role of task-positive control and somatomotor networks for sentence processing with increased semantic integration demands. Additionally, stronger interactions between various task-positive as well as default mode networks were associated with more efficient processing during task conditions with increased semantic integration demands. Collectively, our findings elucidate the complex network interactions underlying semantic integration, a language ability that is crucial for everyday communication. By including young, middle-aged and old adults, our findings provide first insight into semantic integration processes across the aging continuum. These results may inform future studies with healthy old as well as clinical populations.

## Acknowledgements

The authors would like to thank Mareike Thomas, Esther Zwicky, Andrea Matusche, Romy Breuer and Vivien Fritsche for their support with neuropsychological testing. Additionally, thanks to Tobias Hopf for his assistance in evaluating the neuropsychological tests. Andreas Deistung is appreciated for his support in collecting the fMRI data.

## Ethics statement

All experimental procedures were approved by the local ethics committee of the University Hospital Halle (Saale) (reference number: 2020-073).

## Patient consent

All participants gave written informed consent.

## Data availability statement

All data are available from the corresponding authors upon reasonable request.

## Statement

The authors declare no potential conflict of interest. All experimental procedures were approved by the local ethics committee of the University Hospital Halle (Saale) (reference number: 2020-073). All participants gave written informed consent. All data are available from the corresponding authors upon reasonable request.

## References

Alain, C., Reinke, K., Mcdonald, K., Chau, W., Tam, F., Pacurar, A., & Graham, S. (2005). Left thalamo-cortical network implicated in successful speech separation and identification. NeuroImage, 26(2), 592–599. 10.1016/j.neuroimage.2005.02.006

Assaf, M., Calhoun, V. D., Kuzu, C. H., Kraut, M. A., Rivkin, P. R., Hart, J., & Pearlson, G. D. (2006). Neural correlates of the object-recall process in semantic memory. Psychiatry Research: Neuroimaging, 147(2–3), 115–126. 10.1016/j.pscychresns.2006.01.002

Badre, D., Poldrack, R. A., Paré-Blagoev, E. J., Insler, R. Z., & Wagner, A. D. (2005). Dissociable Controlled Retrieval and Generalized Selection Mechanisms in Ventrolateral Prefrontal Cortex. Neuron, 47(6), 907–918. 10.1016/j.neuron.2005.07.023

Badre, D., & Wagner, A. D. (2002). Semantic Retrieval, Mnemonic Control, and Prefrontal Cortex. Behavioral and Cognitive Neuroscience Reviews, 1(3), 206–218. 10.1177/1534582302001003002

Badre, D., & Wagner, A. D. (2007). Left ventrolateral prefrontal cortex and the cognitive control of memory. Neuropsychologia, 45(13), 2883–2901. 10.1016/j.neuropsychologia.2007.06.015

Bates, D., Mächler, M., Bolker, B., & Walker, S. (2014). Fitting Linear Mixed-Effects Models using lme4 (arXiv:1406.5823). arXiv. http://arxiv.org/abs/1406.5823

Baumgaertner, A., Weiller, C., & Büchel, C. (2002). Event-Related fMRI Reveals Cortical Sites Involved in Contextual Sentence Integration. NeuroImage, 16(3), 736–745. 10.1006/nimg.2002.1134

Beck, A. T., Steer, R. A., Ball, R., & Ranieri, W. F. (1996). Comparison of Beck Depression Inventories-IA and-II in Psychiatric Outpatients. J Pers Assess, 67, 588–597.

Brett, M., Anton, J.-L., Valabregue, R., & Poline, J.-B. (2002). Region of interest analysis using an SPM toolbox. NeuroImage, 16(02).

Brockett, A. T., & Roesch, M. R. (2021). The ever-changing OFC landscape: What neural signals in OFC can tell us about inhibitory control. Behavioral Neuroscience, 135(2), 129–137. 10.1037/bne0000412

Calhoun, V. D., Adali, T., Pearlson, G. D., & Pekar, J. J. (2001). A method for making group inferences from functional MRI data using independent component analysis. Human Brain Mapping, 14(3), 140–151. 10.1002/hbm.1048

Calhoun, V. D., Adali, T., Pearlson, G. D., & Pekar, J. J. (2002). ON COMPLEX INFOMAX APPLIED TO FUNCTIONAL MRI DATA. *EEE International Conference on Acoustics*, Speech, and Signal Processing, 1, I–1009.

Calhoun, V. D., Liu, J., & Adalı, T. (2009). A review of group ICA for fMRI data and ICA for joint inference of imaging, genetic, and ERP data. NeuroImage, 45(1), S163–S172. 10.1016/j.neuroimage.2008.10.057

Chan, M. Y., Park, D. C., Savalia, N. K., Petersen, S. E., & Wig, G. S. (2014). Decreased segregation of brain systems across the healthy adult lifespan. Proceedings of the National Academy of Sciences, 111(46). 10.1073/pnas.1415122111

Cole, M. W., Reynolds, J. R., Power, J. D., Repovs, G., Anticevic, A., & Braver, T. S. (2013). Multi-task connectivity reveals flexible hubs for adaptive task control. Nature Neuroscience, 16(9), 1348–1355. 10.1038/nn.3470

Corbetta, M., & Shulman, G. L. (2002). Control of goal-directed and stimulus-driven attention in the brain. Nature Reviews Neuroscience, 3(3), 201–215. 10.1038/nrn755

Davey, J., Thompson, H. E., Hallam, G., Karapanagiotidis, T., Murphy, C., De Caso, I., Krieger-Redwood, K., Bernhardt, B. C., Smallwood, J., & Jefferies, E. (2016). Exploring the role of the posterior middle temporal gyrus in semantic cognition: Integration of anterior temporal lobe with executive processes. NeuroImage, 137, 165–177. 10.1016/j.neuroimage.2016.05.051

DeLong, K. A., Quante, L., & Kutas, M. (2014). Predictability, plausibility, and two late ERP positivities during written sentence comprehension. Neuropsychologia, 61, 150–162. 10.1016/j.neuropsychologia.2014.06.016

Dobbins, I. G., & Wagner, A. D. (2005). Domain-general and Domain-sensitive Prefrontal Mechanisms for Recollecting Events and Detecting Novelty. Cerebral Cortex, 15(11), 1768– 1778. 10.1093/cercor/bhi054

Dosenbach, N. U. F., Fair, D. A., Miezin, F. M., Cohen, A. L., Wenger, K. K., Dosenbach, R. A. T., Fox, M. D., Snyder, A. Z., Vincent, J. L., Raichle, M. E., Schlaggar, B. L., & Petersen, S. E. (2007). Distinct brain networks for adaptive and stable task control in humans. Proceedings of the National Academy of Sciences, 104(26), 11073–11078. 10.1073/pnas.0704320104

Duncan, J. (2010). The multiple-demand (MD) system of the primate brain: Mental programs for intelligent behaviour. Trends in Cognitive Sciences, 14(4), 172–179. 10.1016/j.tics.2010.01.004

Eickhoff, S. B., Stephan, K. E., Mohlberg, H., Grefkes, C., Fink, G. R., Amunts, K., & Zilles, K. (2005). A new SPM toolbox for combining probabilistic cytoarchitectonic maps and functional imaging data. NeuroImage, 25(4), 1325–1335. 10.1016/j.neuroimage.2004.12.034

Esteban, O., Markiewicz, C. J., Blair, R. W., Moodie, C. A., Isik, A. I., Erramuzpe, A., Kent, J. D., Goncalves, M., DuPre, E., Snyder, M., Oya, H., Ghosh, S. S., Wright, J., Durnez, J., Poldrack, R. A., & Gorgolewski, K. J. (2019). fMRIPrep: A robust preprocessing pipeline for functional MRI. Nature Methods, 16(1), 111–116. 10.1038/s41592-018-0235-4

Federmeier, K. D. (2007). Thinking ahead: The role and roots of prediction in language comprehension. Psychophysiology, 44(4), 491–505. 10.1111/j.1469-8986.2007.00531.x

Fedorenko, E., Duncan, J., & Kanwisher, N. (2013). Broad domain generality in focal regions of frontal and parietal cortex. Proceedings of the National Academy of Sciences, 110(41), 16616–16621. 10.1073/pnas.1315235110

Fedorenko, E., & Thompson-Schill, S. L. (2014). Reworking the language network. Trends in Cognitive Sciences, 18(3), 120–126. 10.1016/j.tics.2013.12.006

Feis, R. A., Smith, S. M., Filippini, N., Douaud, G., Dopper, E. G. P., Heise, V., Trachtenberg, A. J., Van Swieten, J. C., Van Buchem, M. A., Rombouts, S. A. R. B., & Mackay, C. E. (2015). ICA-based artifact removal diminishes scan site differences in multi-center resting-state fMRI. Frontiers in Neuroscience, 9. 10.3389/fnins.2015.00395

Folstein, M. F., Folstein, S. E., & McHugh, P. R. (1975). “Mini-mental state”. Journal of Psychiatric Research, 12(3), 189–198. 10.1016/0022-3956(75)90026-6

Fornito, A., Harrison, B. J., Zalesky, A., & Simons, J. S. (2012). Competitive and cooperative dynamics of large-scale brain functional networks supporting recollection. Proceedings of the National Academy of Sciences, 109(31), 12788–12793. 10.1073/pnas.1204185109

Fox, M. D., Snyder, A. Z., Vincent, J. L., Corbetta, M., Van Essen, D. C., & Raichle, M. E. (2005). The human brain is intrinsically organized into dynamic, anticorrelated functional networks. Proceedings of the National Academy of Sciences, 102(27), 9673–9678. 10.1073/pnas.0504136102

Franzmeier, I., Hutton, S. B., & Ferstl, E. C. (2012). The role of the temporal lobe in contextual sentence integration: A single-pulse transcranial magnetic stimulation study. Cognitive Neuroscience, 3(1), 1–7. 10.1080/17588928.2011.556248

Friederici, A. D. (2006). What’s in control of language? Nature Neuroscience, 9(8), 991–992. 10.1038/nn0806-991

Fritsch, M., Rangus, I., & Nolte, C. H. (2022). Thalamic Aphasia: A Review. Current Neurology and Neuroscience Reports, 22(12), 855–865. 10.1007/s11910-022-01242-2

Gold, B. T., Balota, D. A., Jones, S. J., Powell, D. K., Smith, C. D., & Andersen, A. H. (2006). Dissociation of Automatic and Strategic Lexical-Semantics: Functional Magnetic Resonance Imaging Evidence for Differing Roles of Multiple Frontotemporal Regions. Journal of Neuroscience, 26(24), 6523–6532. 10.1523/JNEUROSCI.0808-06.2006

Gorgolewski, K., Burns, C. D., Madison, C., Clark, D., Halchenko, Y. O., Waskom, M. L., & Ghosh, S. S. (2011). Nipype: A Flexible, Lightweight and Extensible Neuroimaging Data Processing Framework in Python. Frontiers in Neuroinformatics, 5. 10.3389/fninf.2011.00013

Grady, C. (2012). The cognitive neuroscience of ageing. Nature Reviews Neuroscience, 13(7), 491–505. 10.1038/nrn3256

Greicius, M. D., Krasnow, B., Reiss, A. L., & Menon, V. (2003). Functional connectivity in the resting brain: A network analysis of the default mode hypothesis. Proceedings of the National Academy of Sciences, 100(1), 253–258. 10.1073/pnas.0135058100

Griffanti, L., Salimi-Khorshidi, G., Beckmann, C. F., Auerbach, E. J., Douaud, G., Sexton, C. E., Zsoldos, E., Ebmeier, K. P., Filippini, N., Mackay, C. E., Moeller, S., Xu, J., Yacoub, E., Baselli, G., Ugurbil, K., Miller, K. L., & Smith, S. M. (2014). ICA-based artefact removal and accelerated fMRI acquisition for improved resting state network imaging. NeuroImage, 95, 232–247. 10.1016/j.neuroimage.2014.03.034

Hagoort, P. (2006). On Broca, Brain, and Binding. In Y. Grodzinsky & K. Amunts (Eds.), Broca’s Region (pp. 242–253). Oxford University Press. 10.1093/acprof:oso/9780195177640.003.0015

Hagoort, P., & Indefrey, P. (2014). The Neurobiology of Language Beyond Single Words. Annual Review of Neuroscience, 37(1), 347–362. 10.1146/annurev-neuro-071013-013847

Hartwigsen, G., Henseler, I., Stockert, A., Wawrzyniak, M., Wendt, C., Klingbeil, J., Baumgaertner, A., & Saur, D. (2017). Integration demands modulate effective connectivity in a fronto-temporal network for contextual sentence integration. NeuroImage, 147, 812–824. 10.1016/j.neuroimage.2016.08.026

Himberg, J., & Hyvarinen, A. (2003). Icasso: Software for investigating the reliability of ICA estimates by clustering and visualization. 2003 IEEE XIII Workshop on Neural Networks for Signal Processing (IEEE Cat. No.03TH8718), 259–268. 10.1109/NNSP.2003.1318025

Hoffman, P. (2018). An individual differences approach to semantic cognition: Divergent effects of age on representation, retrieval and selection. Scientific Reports, 8(1), 8145. 10.1038/s41598-018-26569-0

Hoffman, P., Cox, S. R., Dykiert, D., Muñoz Maniega, S., Valdés Hernández, M. C., Bastin, M. E., Wardlaw, J. M., & Deary, I. J. (2017). Brain grey and white matter predictors of verbal ability traits in older age: The Lothian Birth Cohort 1936. NeuroImage, 156, 394–402. 10.1016/j.neuroimage.2017.05.052

Hoffman, P., & MacPherson, S. E. (2022). What determines cognitive estimation ability? Changing contributions of semantic and executive domains as a function of age. Journal of Neuropsychology, 16(3), 481–497. 10.1111/jnp.12279

Hoffman, P., McClelland, J. L., & Lambon Ralph, M. A. (2018). Concepts, control, and context: A connectionist account of normal and disordered semantic cognition. Psychological Review, 125(3), 293–328. 10.1037/rev0000094

Humphreys, G. F., Hoffman, P., Visser, M., Binney, R. J., & Lambon Ralph, M. A. (2015). Establishing task- and modality-dependent dissociations between the semantic and default mode networks. Proceedings of the National Academy of Sciences, 112(25), 7857–7862. 10.1073/pnas.1422760112

Humphreys, G. F., Jackson, R. L., & Lambon Ralph, M. A. (2019). Overarching principles and dimensions of the functional organisation in the inferior parietal cortex. 10.1101/654178

Humphreys, G. F., & Lambon Ralph, M. A. (2015). Fusion and Fission of Cognitive Functions in the Human Parietal Cortex. Cerebral Cortex, 25(10), 3547–3560. 10.1093/cercor/bhu198

Humphries, C., Binder, J. R., Medler, D. A., & Liebenthal, E. (2007). Time course of semantic processes during sentence comprehension: An fMRI study. NeuroImage, 36(3), 924–932. 10.1016/j.neuroimage.2007.03.059

Hwang, K., Bertolero, M. A., Liu, W. B., & D’Esposito, M. (2017). The Human Thalamus Is an Integrative Hub for Functional Brain Networks. The Journal of Neuroscience, 37(23), 5594– 5607. 10.1523/JNEUROSCI.0067-17.2017

Jaccard, P. (1912). THE DISTRIBUTION OF THE FLORA IN THE ALPINE ZONE.1. New Phytologist, 11(2), 37–50. 10.1111/j.1469-8137.1912.tb05611.x

Jackson, R. L. (2021). The neural correlates of semantic control revisited. NeuroImage, 224, 117444. 10.1016/j.neuroimage.2020.117444

Jackson, R. L., Hoffman, P., Pobric, G., & Lambon Ralph, M. A. (2016). The Semantic Network at Work and Rest: Differential Connectivity of Anterior Temporal Lobe Subregions. The Journal of Neuroscience, 36(5), 1490–1501. 10.1523/JNEUROSCI.2999-15.2016

Jefferies, E. (2013). The neural basis of semantic cognition: Converging evidence from neuropsychology, neuroimaging and TMS. Cortex, 49(3), 611–625. 10.1016/j.cortex.2012.10.008

Jefferies, E., & Lambon Ralph, M. A. (2006). Semantic impairment in stroke aphasia versus semantic dementia: A case-series comparison. Brain, 129(8), 2132–2147. 10.1093/brain/awl153

Jung, J., Cloutman, L. L., Binney, R. J., & Lambon Ralph, M. A. (2017). The structural connectivity of higher order association cortices reflects human functional brain networks. Cortex, 97, 221–239. 10.1016/j.cortex.2016.08.011

Ketteler, D., Kastrau, F., Vohn, R., & Huber, W. (2008). The subcortical role of language processing. High level linguistic features such as ambiguity-resolution and the human brain; an fMRI study. NeuroImage, 39(4), 2002–2009. 10.1016/j.neuroimage.2007.10.023

Kielar, A., Deschamps, T., Jokel, R., & Meltzer, J. A. (2016). Functional reorganization of language networks for semantics and syntax in chronic stroke: Evidence from MEG. Human Brain Mapping, 37(8), 2869–2893. 10.1002/hbm.23212

Kotz, S. (2002). Modulation of the Lexical–Semantic Network by Auditory Semantic Priming: An Event-Related Functional MRI Study. NeuroImage, 17(4), 1761–1772. 10.1006/nimg.2002.1316

Krieger-Redwood, K., Jefferies, E., Karapanagiotidis, T., Seymour, R., Nunes, A., Ang, J. W. A., Majernikova, V., Mollo, G., & Smallwood, J. (2016). Down but not out in posterior cingulate cortex: Deactivation yet functional coupling with prefrontal cortex during demanding semantic cognition. NeuroImage, 141, 366–377. 10.1016/j.neuroimage.2016.07.060

Krieger-Redwood, K., Teige, C., Davey, J., Hymers, M., & Jefferies, E. (2015). Conceptual control across modalities: Graded specialisation for pictures and words in inferior frontal and posterior temporal cortex. Neuropsychologia, 76, 92–107. 10.1016/j.neuropsychologia.2015.02.030

Kutas, M., & Federmeier, K. D. (2000). Electrophysiology reveals semantic memory use in language comprehension. Trends in Cognitive Sciences, 4(12), 463–470. 10.1016/S1364-6613(00)01560-6

Kutas, M., & Hillyard, S. A. (1980). Reading senseless sentences: Brain potentials reflect semantic incongruity. *Science*, New Series(Vol. 207, No. 4427), 203–205.

Lambon Ralph, M. A., Jefferies, E., Patterson, K., & Rogers, T. T. (2017). The neural and computational bases of semantic cognition. Nature Reviews Neuroscience, 18(1), 42–55. 10.1038/nrn.2016.150

Lanzoni, L., Ravasio, D., Thompson, H., Vatansever, D., Margulies, D., Smallwood, J., & Jefferies, E. (2020). The role of default mode network in semantic cue integration. NeuroImage, 219, 117019. 10.1016/j.neuroimage.2020.117019

Lau, E. F., Phillips, C., & Poeppel, D. (2008). A cortical network for semantics: (De)constructing the N400. Nature Reviews Neuroscience, 9(12), 920–933. 10.1038/nrn2532

Lenth, R. (2020). Emmeans: Estimated marginal means, aka leastsquares means. R Package Version 1.4.8.

Lüdecke, D. (2018). ggeffects: Tidy Data Frames of Marginal Effects from Regression Models. Journal of Open Source Software, 3(26), 772. 10.21105/joss.00772

Marek, S., & Dosenbach, N. U. F. (2018). The frontoparietal network: Function, electrophysiology, and importance of individual precision mapping. Dialogues in Clinical Neuroscience, 20(2), 133–140. 10.31887/DCNS.2018.20.2/smarek

Martin, S., Saur, D., & Hartwigsen, G. (2022). Age-Dependent Contribution of Domain-General Networks to Semantic Cognition. Cerebral Cortex, 32(4), 870–890. 10.1093/cercor/bhab252

Martin, S., Williams, K. A., Saur, D., & Hartwigsen, G. (2023). Age-related reorganization of functional network architecture in semantic cognition. Cerebral Cortex, 33(8), 4886–4903. 10.1093/cercor/bhac387

Mascali, D., DiNuzzo, M., Serra, L., Mangia, S., Maraviglia, B., Bozzali, M., & Giove, F. (2018). Disruption of Semantic Network in Mild Alzheimer’s Disease Revealed by Resting-State fMRI. Neuroscience, 371, 38–48. 10.1016/j.neuroscience.2017.11.030

McLaren, D. G., Ries, M. L., Xu, G., & Johnson, S. C. (2012). A generalized form of context-dependent psychophysiological interactions (gPPI): A comparison to standard approaches. NeuroImage, 61(4), 1277–1286. 10.1016/j.neuroimage.2012.03.068

Menon, V., & D’Esposito, M. (2022). The role of PFC networks in cognitive control and executive function. Neuropsychopharmacology, 47(1), 90–103. 10.1038/s41386-021-01152-w

Mion, M., Patterson, K., Acosta-Cabronero, J., Pengas, G., Izquierdo-Garcia, D., Hong, Y. T., Fryer, T. D., Williams, G. B., Hodges, J. R., & Nestor, P. J. (2010). What the left and right anterior fusiform gyri tell us about semantic memory. Brain, 133(11), 3256–3268. 10.1093/brain/awq272

Morcom, A. M., & Johnson, W. (2015). Neural Reorganization and Compensation in Aging. Journal of Cognitive Neuroscience, 27(7), 1275–1285. 10.1162/jocn_a_00783

Nadeau, S., & Crosson, B. (1997). Subcortical aphasia. Brain and Language, 58(3), 355–402.

Nagel, I. E., Schumacher, E. H., Goebel, R., & D’Esposito, M. (2008). Functional MRI investigation of verbal selection mechanisms in lateral prefrontal cortex. NeuroImage, 43(4), 801–807. 10.1016/j.neuroimage.2008.07.017

Noonan, K. A., Jefferies, E., Visser, M., & Lambon Ralph, M. A. (2013). Going beyond Inferior Prefrontal Involvement in Semantic Control: Evidence for the Additional Contribution of Dorsal Angular Gyrus and Posterior Middle Temporal Cortex. Journal of Cognitive Neuroscience, 25(11), 1824–1850. 10.1162/jocn_a_00442

Obleser, J., & Kotz, S. A. (2010). Expectancy Constraints in Degraded Speech Modulate the Language Comprehension Network. Cerebral Cortex, 20(3), 633–640. 10.1093/cercor/bhp128

Oldfield, R. (1971). The assessment and analysis of handeness: The Edinburgh inventory. Neuropsychologia, 9, 97–113.

Patterson, K., Nestor, P. J., & Rogers, T. T. (2007). Where do you know what you know? The representation of semantic knowledge in the human brain. Nature Reviews Neuroscience, 8(12), 976–987. 10.1038/nrn2277

Peirce, J., Gray, J. R., Simpson, S., MacAskill, M., Höchenberger, R., Sogo, H., Kastman, E., & Lindeløv, J. K. (2019). PsychoPy2: Experiments in behavior made easy. Behavior Research Methods, 51(1), 195–203. 10.3758/s13428-018-01193-y

Pobric, G., Jefferies, E., & Ralph, M. A. L. (2007). Anterior temporal lobes mediate semantic representation: Mimicking semantic dementia by using rTMS in normal participants. Proceedings of the National Academy of Sciences, 104(50), 20137–20141. 10.1073/pnas.0707383104

Power, J. D., Cohen, A. L., Nelson, S. M., Wig, G. S., Barnes, K. A., Church, J. A., Vogel, A. C., Laumann, T. O., Miezin, F. M., Schlaggar, B. L., & Petersen, S. E. (2011). Functional Network Organization of the Human Brain. Neuron, 72(4), 665–678. 10.1016/j.neuron.2011.09.006

Price, A. R., Peelle, J. E., Bonner, M. F., Grossman, M., & Hamilton, R. H. (2016). Causal Evidence for a Mechanism of Semantic Integration in the Angular Gyrus as Revealed by High-Definition Transcranial Direct Current Stimulation. The Journal of Neuroscience, 36(13), 3829–3838. 10.1523/JNEUROSCI.3120-15.2016

Price, C. J., Moore, C. J., Humphreys, G. W., & Wise, R. J. S. (1997). Segregating Semantic from Phonological Processes during Reading. Journal of Cognitive Neuroscience, *Vol.* 9*, No.* *6*, p.727–733.

Qin, P., & Northoff, G. (2011). How is our self related to midline regions and the default-mode network? NeuroImage, 57(3), 1221–1233. 10.1016/j.neuroimage.2011.05.028

R Core Team. (2021). R: A language and environment for statistical computing. Vienna, *Austria*: *R Foundation for Statistical Computing*.

Radanovic, M., & Almeida, V. N. (2021). Subcortical Aphasia. Current Neurology and Neuroscience Reports, 21(12), 73. 10.1007/s11910-021-01156-5

Rissanen, J. (1978). Modeling by shortest data description. Automatica, 14(5), 465–471. 10.1016/0005-1098(78)90005-5

Rodd, J. M., Davis, M. H., & Johnsrude, I. S. (2005). The Neural Mechanisms of Speech Comprehension: fMRI studies of Semantic Ambiguity. Cerebral Cortex, 15(8), 1261–1269. 10.1093/cercor/bhi009

Rogalsky, C., & Hickok, G. (2009). Selective Attention to Semantic and Syntactic Features Modulates Sentence Processing Networks in Anterior Temporal Cortex. Cerebral Cortex, 19(4), 786–796. 10.1093/cercor/bhn126

Ryskin, R., & Nieuwland, M. S. (2023). Prediction during language comprehension: What is next? Trends in Cognitive Sciences, 27(11), 1032–1052. 10.1016/j.tics.2023.08.003

Schwartz, G. (1978). Estimating the dimension of a model. The Annals of Statistics, 6.

Seghier, M. L. (2013). The Angular Gyrus: Multiple Functions and Multiple Subdivisions. The Neuroscientist, 19(1), 43–61. 10.1177/1073858412440596

Setton, R., Mwilambwe-Tshilobo, L., Girn, M., Lockrow, A. W., Baracchini, G., Hughes, C., Lowe, A. J., Cassidy, B. N., Li, J., Luh, W.-M., Bzdok, D., Leahy, R. M., Ge, T., Margulies, D. S., Misic, B., Bernhardt, B. C., Stevens, W. D., De Brigard, F., Kundu, P., … Spreng, R. N. (2022). Age differences in the functional architecture of the human brain. Cerebral Cortex, 33(1), 114–134. 10.1093/cercor/bhac056

Smallwood, J., Bernhardt, B. C., Leech, R., Bzdok, D., Jefferies, E., & Margulies, D. S. (2021). The default mode network in cognition: A topographical perspective. Nature Reviews Neuroscience, 22(8), 503–513. 10.1038/s41583-021-00474-4

Spreng, R. N., Stevens, W. D., Chamberlain, J. P., Gilmore, A. W., & Schacter, D. L. (2010). Default network activity, coupled with the frontoparietal control network, supports goal-directed cognition. NeuroImage, 53(1), 303–317. 10.1016/j.neuroimage.2010.06.016

Thompson-Schill, S. L., D’Esposito, M., Aguirre, G. K., & Farah, M. J. (1997). Role of left inferior prefrontal cortex in retrieval of semantic knowledge: A reevaluation. Proceedings of the National Academy of Sciences, 94(26), 14792–14797. 10.1073/pnas.94.26.14792

Turker, S., Kuhnke, P., Eickhoff, S. B., Caspers, S., & Hartwigsen, G. (2023). Cortical, subcortical, and cerebellar contributions to language processing: A meta-analytic review of 403 neuroimaging experiments. Psychological Bulletin, 149(11–12), 699–723. 10.1037/bul0000403

Von Der Heide, R. J., Skipper, L. M., Klobusicky, E., & Olson, I. R. (2013). Dissecting the uncinate fasciculus: Disorders, controversies and a hypothesis. Brain, 136(6), 1692–1707. 10.1093/brain/awt094

Wawrzyniak, M., Hoffstaedter, F., Klingbeil, J., Stockert, A., Wrede, K., Hartwigsen, G., Eickhoff, S. B., Classen, J., & Saur, D. (2017). Fronto-temporal interactions are functionally relevant for semantic control in language processing. PLOS ONE, 12(5), e0177753. 10.1371/journal.pone.0177753

Wechsler, D., & De Lemos, M. M. (1981). Wechsler adult intelligence scale-revised. *New York*: Psychological Corporation, 1.

Whitney, C., Kirk, M., O’Sullivan, J., Lambon Ralph, M. A., & Jefferies, E. (2011). The Neural Organization of Semantic Control: TMS Evidence for a Distributed Network in Left Inferior Frontal and Posterior Middle Temporal Gyrus. Cerebral Cortex, 21(5), 1066–1075. 10.1093/cercor/bhq180

Whitney, C., Kirk, M., O’Sullivan, J., Lambon Ralph, M. A., & Jefferies, E. (2012). Executive Semantic Processing Is Underpinned by a Large-scale Neural Network: Revealing the Contribution of Left Prefrontal, Posterior Temporal, and Parietal Cortex to Controlled Retrieval and Selection Using TMS. Journal of Cognitive Neuroscience, 24(1), 133–147. 10.1162/jocn_a_00123

Wickham, H. (2016). Ggplot2. Elegant Graphics for Data Analysis, Use R! Cham, *Switzerland*: *Springer International Publishing*.

Wu, W., & Hoffman, P. (2022). Validated measures of semantic knowledge and semantic control: Normative data from young and older adults for more than 300 semantic judgements. Royal Society Open Science.

Yee, E., & Thompson-Schill, S. L. (2016). Putting concepts into context. Psychonomic Bulletin & Review, 23(4), 1015–1027. 10.3758/s13423-015-0948-7

Yeo, B. T. T., Krienen, F. M., Sepulcre, J., Sabuncu, M. R., Lashkari, D., Hollinshead, M., Roffman, J. L., Smoller, J. W., Zöllei, L., Polimeni, J. R., Fischl, B., Liu, H., & Buckner, R. L. (2011). The organization of the human cerebral cortex estimated by intrinsic functional connectivity. Journal of Neurophysiology, 106(3), 1125–1165. 10.1152/jn.00338.2011

Zalesky, A., Fornito, A., & Bullmore, E. T. (2010). Network-based statistic: Identifying differences in brain networks. NeuroImage, 53(4), 1197–1207. 10.1016/j.neuroimage.2010.06.041

Zhu, Z., Feng, G., Zhang, J. X., Li, G., Li, H., & Wang, S. (2013). The role of the left prefrontal cortex in sentence-level semantic integration. NeuroImage, 76, 325–331. 10.1016/j.neuroimage.2013.02.060

Zimmermann, P., & Fimm, B. (2002). A test battery for attentional performance. *Applied Neuropsychology of Attention. Theory*, Diagnosis and Rehabilitation., 110, 151.

