## supplemental material for "Semantic integration demands modulate large-scale network interactions in the brain"

### Supplementary Methods

**Figure S1**

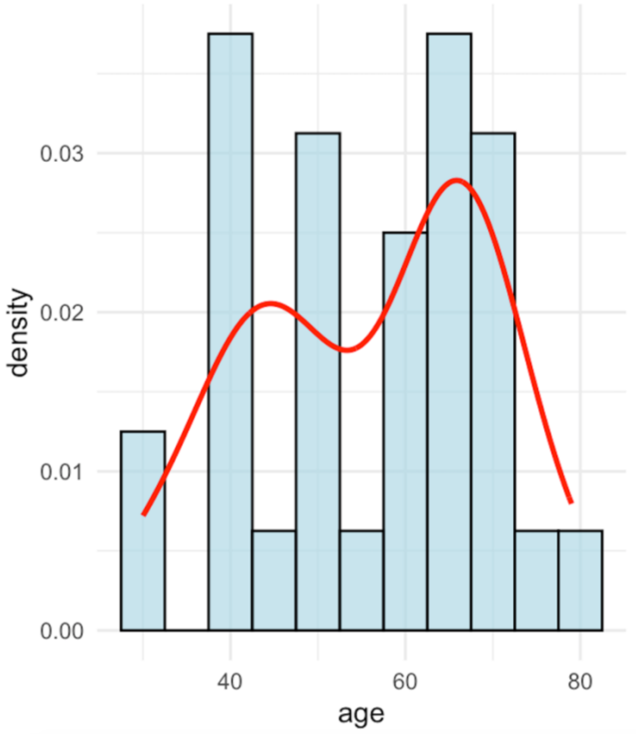

**Figure S1:** *Age distribution of participants.* The density curve (red) represents the estimated probability density function of the age distribution.

**Table S1.** *Demographic and neuropsychological test characteristics*

| | Mean values of raw scores (SD) |
|---|---|
| **Demographics** | |
| **Age (years)** | 55 (12.77) |
| **Sex (F:M)** | 15:17 |
| **Education (years)** | 16 (2.44) |
| **Laterality Quotient** | 91.48 (9) |
| **Neuropsychological Test** | |
| **MMSE** | 29.281 (1.05) |
| **BDI-II** | 3.594 (3.38) |
| **Digit span forward** | 6.5 (1.14) |
| **Digit span backward** | 5.188 (0.99) |
| **TAP (sum score)** | 193.84 |

***Abbreviations:*** *BDI-II: Beck Depression Inventory; F: Female; M: Male; MMSE: Mini Mental State Examination; SD: Standard Deviation; TAP: Test of Attentional Performance.*

**Equation S1.** *Linear mixed-effects model for analysis of reaction times*

log (Reaction time) = β0 + β1condition + β2age + β3education + β4condition x age + (1|subject) + (1|stimulus) + ε

**Equation S2.** *Generalized linear mixed-effects logistic regression for analysis of accuracy*

Error rate = β0 + β1condition + β2age + β3education + (1|subject) + (1|stimulus) + ε

**Equation S3.** *Linear mixed-effects model for analysis of cPPI connectivity effects on reaction times*

log (Reaction time) = β0 + β1network measure + β2condition + β3age + βeducation +β4network measure x condition + (1|subject) +ε

**Equation S4.** *Generalized linear mixed-effects logistic regression for analysis of cPPI connectivity effects on accuracy*

Error rate = β0 + β1network measure + β2condition + β3age + β4network measure x condition + (1|subject) + ε

#### **Note S1.** *Preprocessing of fMRI-data using fMRIPrep*

Results included in this manuscript come from preprocessing performed using *fMRIPrep* 23.0.0 (Esteban, Markiewicz, et al. (2018); Esteban, Blair, et al. (2018); RRID:SCR_016216), which is based on *Nipype* 1.8.5 (K. Gorgolewski et al. (2011); K. J. Gorgolewski et al. (2018); RRID:SCR_002502)

###### Anatomical data preprocessing

A total of 1 T1-weighted (T1w) images were found within the input BIDS dataset.The T1-weighted (T1w) image was corrected for intensity non-uniformity (INU) with N4BiasFieldCorrection (Tustison et al. 2010), distributed with ANTs 2.3.3 (Avants et al. 2008, RRID:SCR_004757), and used as T1w-reference throughout the workflow. The T1w-reference was then skull-stripped with a *Nipype* implementation of the antsBrainExtraction.sh workflow (from ANTs), using OASIS30ANTs as target template. Brain tissue segmentation of cerebrospinal fluid (CSF), white-matter (WM) and gray-matter (GM) was performed on the brain-extracted T1w using fast (FSL 6.0.5.1:57b01774, RRID:SCR_002823, Zhang, Brady, and Smith 2001). Brain surfaces were reconstructed using recon-all (FreeSurfer 7.3.2, RRID:SCR_001847, Dale, Fischl, and Sereno 1999), and the brain mask estimated previously was refined with a custom variation of the method to reconcile ANTs-derived and FreeSurfer-derived segmentations of the cortical gray-matter of Mindboggle (RRID:SCR_002438, Klein et al. 2017). Volume-based spatial normalization to two standard spaces (MNI152NLin6Asym, MNI152NLin2009cAsym) was performed through nonlinear registration with antsRegistration (ANTs 2.3.3), using brain-extracted versions of both T1w reference and the T1w template. The following templates were selected for spatial normalization: *FSL's MNI ICBM 152 non-linear 6th Generation Asymmetric Average Brain Stereotaxic Registration Model* [Evans et al. (2012), RRID:SCR_002823; TemplateFlow ID: MNI152NLin6Asym], *ICBM 152 Nonlinear Asymmetrical template version 2009c* [Fonov et al. (2009), RRID:SCR_008796; TemplateFlow ID: MNI152NLin2009cAsym].

###### Functional data preprocessing

For each of the 5 BOLD runs found per subject (across all tasks and sessions), the following preprocessing was performed. First, a reference volume and its skull-stripped version were generated using a custom methodology of *fMRIPrep*. Head-motion parameters with respect to the BOLD reference (transformation matrices, and six corresponding rotation and translation parameters) are estimated before any spatiotemporal filtering using mcflirt (FSL 6.0.5.1:57b01774, Jenkinson et al. 2002). BOLD runs were slice-time corrected to 0.962s (0.5 of slice acquisition range 0s-1.92s) using 3dTshift from AFNI (Cox and Hyde 1997, RRID:SCR_005927). The BOLD time-series (including slice-timing correction when applied) were resampled onto their original, native space by applying the transforms to correct for head-motion. These resampled BOLD time-series will be referred to as *preprocessed BOLD in original space*, or just *preprocessed BOLD*. The BOLD reference was then co-registered to the T1w reference using bbregister (FreeSurfer) which implements boundary-based registration (Greve and Fischl 2009). Co-registration was configured with six degrees of freedom. Several confounding time-series were calculated based on the *preprocessed BOLD*: framewise displacement (FD), DVARS and three region-wise global signals. FD was computed using two formulations following Power (absolute sum of relative motions, Power et al. (2014)) and Jenkinson (relative root mean square displacement between affines, Jenkinson et al. (2002)). FD and DVARS are calculated for each functional run, both using their implementations in *Nipype* (following the definitions by Power et al. 2014). The three global signals are extracted within the CSF, the WM, and the whole-brain masks. Additionally, a set of physiological regressors were extracted to allow for component-based noise correction (*CompCor*, Behzadi et al. 2007). Principal components are estimated after high-pass filtering the *preprocessed BOLD* time-series (using a discrete cosine filter with 128s cut-off) for the two *CompCor* variants: temporal (tCompCor) and anatomical (aCompCor). tCompCor components are then calculated from the top 2% variable voxels within the brain mask. For aCompCor, three probabilistic masks (CSF, WM and combined CSF+WM) are generated in anatomical space. The implementation differs from that of Behzadi et al. in that instead of eroding the masks by 2 pixels on BOLD space, a mask of pixels that likely contain a volume fraction of GM is subtracted from the aCompCor masks. This mask is obtained by dilating a GM mask extracted from the FreeSurfer's *aseg* segmentation, and it ensures components are not extracted from voxels containing a minimal fraction of GM. Finally, these masks are resampled into BOLD space and binarized by thresholding at 0.99 (as in the original implementation). Components are also calculated separately within the WM and CSF masks. For each CompCor decomposition, the *k* components with the largest singular values are retained, such that the retained components' time series are sufficient to explain 50 percent of variance across the nuisance mask (CSF, WM, combined, or temporal). The remaining components are dropped from consideration. The head-motion estimates calculated in the correction step were also placed within the corresponding confounds file. The confound time series derived from head motion estimates and global signals were expanded with the inclusion of temporal derivatives and quadratic terms for each (Satterthwaite et al. 2013). Frames that exceeded a threshold of 0.7 mm FD or 1.5 standardized DVARS were annotated as motion outliers. Additional nuisance timeseries are calculated by means of principal components analysis of the signal found within a thin band (*crown*) of voxels around the edge of the brain, as proposed by (Patriat, Reynolds, and Birn 2017). The BOLD time-series were resampled into standard space, generating a *preprocessed BOLD run in MNI152NLin6Asym space*. First, a reference volume and its skull-stripped version were generated using a custom methodology of *fMRIPrep*. All resamplings can be performed with *a single interpolation step* by composing all the pertinent transformations (i.e. head-motion transform matrices, susceptibility distortion correction when available, and co-registrations to anatomical and output spaces). Gridded (volumetric) resamplings were performed using antsApplyTransforms (ANTs), configured with Lanczos interpolation to minimize the smoothing effects of other kernels (Lanczos 1964). Non-gridded (surface) resamplings were performed using mri_vol2surf (FreeSurfer).

Many internal operations of *fMRIPrep* use *Nilearn* 0.9.1 (Abraham et al. 2014, RRID:SCR_001362), mostly within the functional processing workflow. For more details of the pipeline, see the section corresponding to workflows in *fMRIPrep*'s documentation.

###### Copyright Waiver

The above boilerplate text was automatically generated by fMRIPrep with the express intention that users should copy and paste this text into their manuscripts *unchanged*. It is released under the CC0 license.

###### Errors

No errors to report!

**Figure S2**

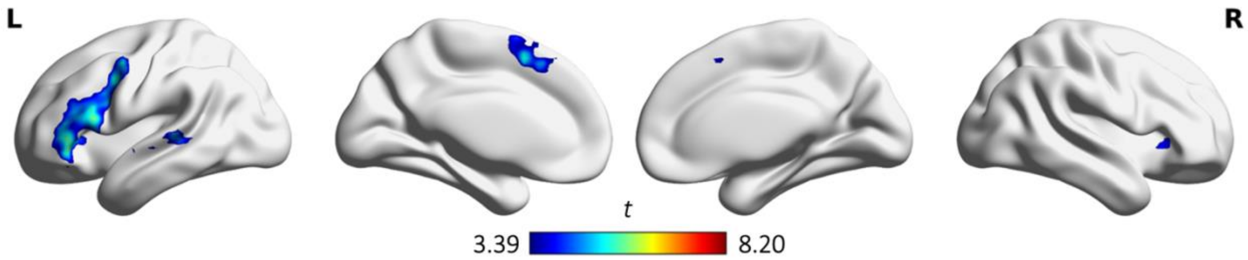

**Figure S2**. *Conjunction of the contrasts "unexpected sentence endings > expected sentence endings ∩ anomalous sentence endings > expected sentence endings"*.
Results are FWE-corrected at $p < 0.05$ at the cluster level.

**Table S2.** *Coordinates of seed regions for gPPI analysis*

| ROI | Region | Side | MNI Coordinates (in mm) | | |
|---|---|---|---|---|---|
| | | | x | y | z |
| **1** | Inferior frontal gyrus (pars triangularis) | L | -54 | 36 | 3 |
| **2** | Presupplementary motor area | L | -6 | 12 | 51 |
| **3** | Cerebellum VI | R | 27 | -66 | -27 |
| **4** | Frontal pole / orbital gyrus | L | 36 | 36 | -15 |
| **5** | Frontal pole / inferior frontal gyrus | R | 48 | 36 | 15 |

***Note:*** *Seed regions were obtained from the conjunction of the contrasts "anomalous > expected" and "unexpected > expected" sentence endings.* ***Abbreviation:*** *ROI: Region of Interest.*

### Supplementary Results

**Table S3.** *Statistical results of mixed-effects models for reaction times and error rates*

| | Log (Reaction times) | | | Error rate | | |
|---|---|---|---|---|---|---|
| ***Predictors*** | *Estimates* | *CI* | *P* | *Odds Ratios* | *CI* | *P* |
| **(Intercept)** | 6.94 | 6.90 – 6.97 | **<0.001** | 0.11 | 0.08 – 0.17 | **<0.001** |
| **condition [unexpected]** | 0.09 | 0.06 – 0.12 | **<0.001** | | | |
| **condition [anomalous]** | 0.24 | 0.21 – 0.27 | **<0.001** | | | |
| **condition [pseudoword]** | 0.26 | 0.24 – 0.29 | **<0.001** | | | |
| **age (centered)** | 0.00 | -0.00 – 0.00 | 0.222 | 1.02 | 1.01 – 1.04 | **0.002** |
| **education (centered)** | 0.00 | -0.01 – 0.02 | 0.736 | 0.98 | 0.90 – 1.07 | 0.645 |
| **condition [unexpected] × age (centered)** | 0.00 | 0.00 – 0.00 | **0.002** | | | |
| **condition [anomalous] × age (centered)** | 0.00 | 0.00 – 0.00 | **0.002** | | | |
| **condition [pseudoword] × age (centered)** | -0.00 | -0.00 – 0.00 | 0.843 | | | |
| **condition [expected]** | | | | 0.09 | 0.05 – 0.17 | **<0.001** |
| **condition [pseudoword]** | | | | 0.35 | 0.22 – 0.56 | **<0.001** |
| **condition [unexpected]** | | | | 0.13 | 0.07 – 0.24 | **<0.001** |
| **Random Effects** | | | | | | |
| **$\sigma^2$** | 0.02 | | | 3.29 | | |
| **$\tau_{00}$** | 0.01 $_{stimulus}$ | | | 1.72 $_{stimulus}$ | | |
| | 0.01 $_{subject}$ | | | 0.27 $_{subject}$ | | |
| **ICC** | 0.46 | | | 0.38 | | |
| **N** | 32 $_{subject}$ | | | 32 $_{subject}$ | | |
| | 300 $_{stimulus}$ | | | 300 $_{stimulus}$ | | |
| **Observations** | 8406 | | | 9565 | | |
| **Marginal $R^2$ / Conditional $R^2$** | 0.255 / 0.595 | | | 0.132 / 0.460 | | |

***Note:*** *Significant effects are marked in bold. Contrasts are treatment coded. P-values were obtained via likelihood ratio tests.* ***Abbreviation:*** *CI: Confidence interval.*

**Figure S3**

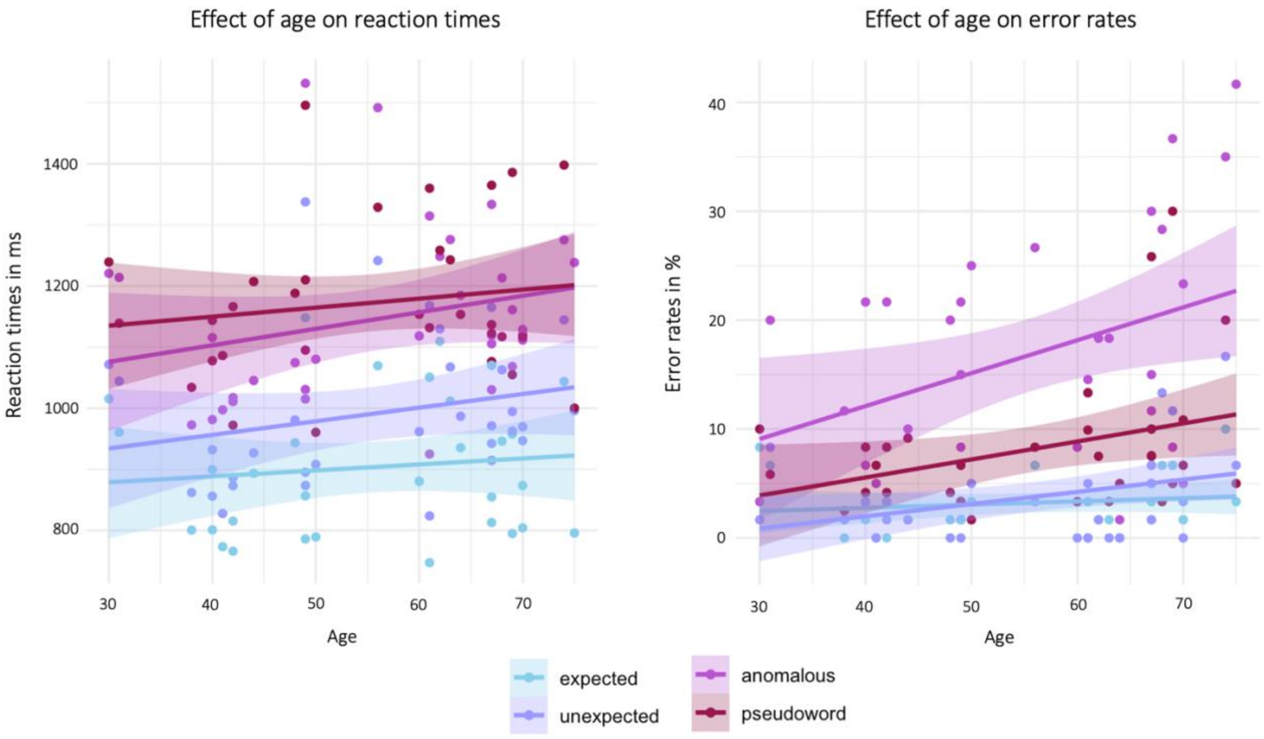

**Figure S3.** *Effect of age on behavioral data.*

**Figure S4**

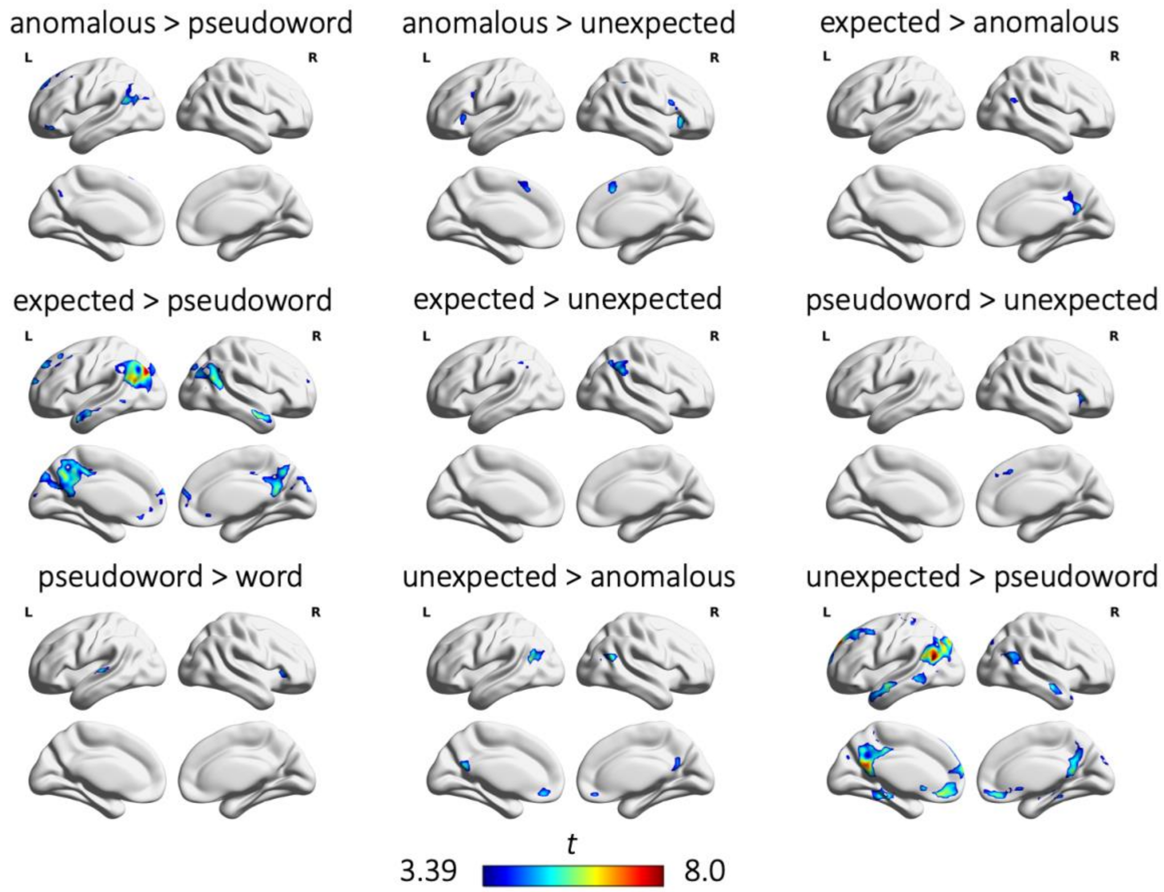

**Figure S4.** *Group-level results from univariate fMRI-analysis. Results are FWE-corrected at p < 0.05 at the cluster level. No significant results were found for the contrast "pseudoword > anomalous".*

**Table S4.** *Group-level activation peaks from univariate fMRI-analysis*

| Region | Side | MNI Coordinates (in mm) | | | t | k |
|---|---|---|---|---|---|---|
| | | x | y | z | | |
| **Anomalous > expected** | | | | | | |
| **Inferior frontal gyrus (p. op.)** | L | -57 | 21 | 24 | 9.15 | 1387 |
| **Presupplementary motor area** | L | -6 | 21 | 45 | 9.06 | 659 |
| **Frontal orbital cortex / insula** | R | 33 | 27 | 0 | 7.96 | 820 |
| **Cerebellum VI** | L | -27 | -66 | -27 | 7.78 | 265 |
| **Superior temporal gyrus** | L | -60 | -30 | 6 | 6.59 | 208 |
| **Cerebellum crus II** | R | 12 | -81 | -30 | 6.07 | 219 |
| **Anomalous > pseudoword** | | | | | | |
| **Frontal pole** | L | -39 | 42 | -15 | 6.16 | 58 |
| **Angular gyrus** | L | -48 | -54 | 24 | 5.88 | 473 |
| **Superior frontal gyrus** | L | -9 | 27 | 63 | 5.32 | 295 |
| **Cerebellum Crus I** | R | 24 | 48 | -24 | 5.08 | 88 |
| **Superior frontal gyrus** | L | -9 | -60 | 33 | 4.60 | 50 |
| **Anomalous > unexpected** | | | | | | |
| **Insula cortex** | R | 33 | 24 | -3 | 5.74 | 96 |
| **Presupplementary motor area** | R | 6 | 21 | 45 | 5.50 | 130 |
| **Inferior parietal lobe** | R | 51 | -42 | 45 | 5.00 | 82 |
| **Inferior frontal gyrus** | R | 45 | 12 | 21 | 4.96 | 87 |
| **Precentral gyrus** | L | -39 | 3 | 33 | 4.88 | 57 |
| **Frontal operculum** | L | -42 | 21 | 6 | 4.82 | 66 |
| **Expected > anomalous** | | | | | | |
| **Precuneous cortex** | R | 9 | -57 | 24 | 5.78 | 189 |
| **Inferior parietal lobe** | R | 42 | -54 | 24 | 4.95 | 75 |
| **Expected > pseudoword** | | | | | | |
| **Inferior parietal lobe** | L | -42 | -78 | 33 | 8.20 | 3387 |
| **Middle temporal gyrus** | R | 60 | -9 | -18 | 6.86 | 187 |
| **Middle frontal gyrus** | L | -39 | 21 | 51 | 6.65 | 357 |
| **Temporal pole** | L | -51 | 6 | -24 | 6.09 | 196 |
| **Frontal pole** | L | -12 | 63 | 21 | 5.94 | 414 |
| **Middle frontal gyrus** | R | 21 | 57 | 39 | 5.75 | 124 |
| **Expected > unexpected** | | | | | | |
| **Angular gyrus** | R | 63 | -48 | 36 | 6.26 | 286 |
| **Supramarginal gyrus** | L | -54 | -48 | 45 | 4.72 | 45 |
| **Pseudoword > expected** | | | | | | |
| **Inferior frontal gyrus (p. op.)** | L | -42 | 6 | 24 | 9.21 | 1163 |
| **Frontal orbital cortex** | R | 33 | 27 | 3 | 7.52 | 193 |
| **Superior temporal gyrus** | R | 63 | -3 | -3 | 7.41 | 89 |
| **Presupplementary motor area** | L | -6 | 12 | 54 | 6.65 | 414 |
| **Middle frontal gyrus** | R | 51 | 36 | 24 | 6.55 | 311 |
| **Pseudoword > unexpected** | | | | | | |
| **Frontal orbital cortex** | R | 39 | 30 | 0 | 5.18 | 64 |
| **Paracingulate gyrus** | R | 6 | 24 | 42 | 4.72 | 44 |
| **Pseudoword > word** | | | | | | |
| **Planum temporale** | L | -63 | -21 | 6 | 5.89 | 72 |
| **Frontal operculum** | R | 36 | 27 | 3 | 4.77 | 62 |

| | | | | | | |
|---|---|---|---|---|---|---|
| **Unexpected > anomalous** | | | | | | |
| **Inferior parietal lobe** | L | -54 | -63 | 18 | 7.19 | 311 |
| **Lateral occipital cortex** | R | 51 | -60 | 21 | 5.91 | 150 |
| **Precuneous cortex** | L | -6 | -57 | 21 | 5.69 | 197 |
| **Frontal medial cortex** | R | 3 | 45 | -15 | 5.26 | 130 |
| **Unexpected > expected** | | | | | | |
| **Inferior frontal gyrus (p. op.)** | L | -39 | 6 | 24 | 7.65 | 1309 |
| **Presupplementary motor area** | L | -3 | 12 | 48 | 5.88 | 237 |
| **Precentral gyrus** | R | 48 | 9 | 27 | 5.16 | 89 |
| **Frontal orbital cortex** | R | 42 | 33 | -12 | 5.07 | 76 |
| **Cerebellum VI** | R | 30 | -66 | -24 | 5.04 | 73 |
| **Inferior temporal gyrus** | L | -54 | -57 | -12 | 4.11 | 42 |
| **Unexpected > pseudoword** | | | | | | |
| **Inferior parietal lobe** | L | -45 | -60 | 18 | 8.70 | 2424 |
| **Frontal pole** | L | -15 | 48 | 39 | 8.34 | 1211 |
| **Middle temporal gyrus** | L | -57 | -9 | -21 | 6.63 | 502 |
| **Middle temporal gyrus** | R | 57 | -9 | -15 | 6.25 | 186 |
| **Inferior parietal lobe** | R | 57 | -60 | 18 | 5.84 | 456 |
| **Middle temporal gyrus** | L | -63 | -42 | -3 | 5.63 | 113 |
| **Parahippocampal gyrus** | R | 30 | -33 | -18 | 5.23 | 107 |
| **Word > pseudoword** | | | | | | |
| **Frontal pole** | L | -15 | 48 | 39 | 11.13 | 1098 |
| **Inferior parietal lobe** | L | -45 | -63 | 21 | 9.41 | 3410 |
| **Middle temporal gyrus** | L | -57 | -9 | -21 | 7.40 | 335 |
| **Parahippocampal gyrus** | L | -30 | -27 | -25 | 6.35 | 122 |
| **Middle temporal gyrus** | R | 57 | 0 | -24 | 6.15 | 215 |

***Note:*** *Cluster corrected at FWE $p < 0.05$ with a voxel-wise threshold at $p < 0.001$. Clusters represent local maxima. Cluster size (k) is given in $mm^3$. No significant cluster were found for the contrast "sentences with pseudoword endings > anomalous endings".* ***Abbreviation:*** *P. op.: Pars opercularis.*

**Table S5.** *Jaccard indices for independent components and cognitive networks* (Yeo et al., 2011)

| | IC04 | IC11 | IC12 | IC13 | IC16 | IC17 | IC18 | IC21 | IC25 | IC33 | IC35 |
|---|---|---|---|---|---|---|---|---|---|---|---|
| **ContA** | 0.002 | 0.024 | 0.050 | 0.031 | 0.011 | 0.075 | 0.019 | 0.001 | ***0.241*** | 0.020 | 0.002 |
| **ContB** | 0.001 | 0.022 | 0.033 | 0.012 | 0.006 | ***0.159*** | 0.004 | 0.005 | 0.064 | 0.033 | 0.050 |
| **ConC** | 0.010 | 0.075 | 0.033 | 0.025 | 0.000 | 0.087 | 0.000 | 0.019 | 0.000 | 0.001 | 0.003 |
| **DefaultA** | 0.011 | ***0.225*** | 0.005 | 0.015 | 0.000 | 0.082 | 0.013 | 0.007 | 0.006 | 0.015 | 0.142 |
| **DefaultB** | 0.026 | 0.023 | 0.034 | 0.014 | 0.005 | 0.048 | 0.004 | 0.007 | 0.081 | 0.078 | ***0.273*** |
| **DefaultC** | 0.000 | 0.059 | 0.000 | 0.017 | 0.014 | 0.002 | 0.016 | 0.045 | 0.000 | 0.002 | 0.003 |
| **DorsAttnA** | 0.000 | 0.056 | 0.083 | 0.040 | **0.035** | 0.013 | 0.112 | 0.051 | 0.064 | 0.000 | 0.002 |
| **DorsAttnB** | 0.004 | 0.018 | *0.154* | 0.079 | 0.010 | 0.005 | 0.016 | 0.001 | 0.013 | 0.001 | 0.002 |
| **LimbicA** | 0.001 | 0.000 | 0.000 | 0.001 | 0.001 | 0.001 | 0.000 | 0.001 | 0.000 | 0.002 | 0.004 |
| **LimbicB** | 0.000 | 0.020 | 0.000 | 0.000 | 0.000 | 0.004 | 0.003 | 0.000 | 0.001 | 0.002 | 0.006 |
| **SalVentAttnA** | 0.087 | 0.040 | 0.063 | ***0.240*** | 0.000 | 0.006 | 0.017 | 0.001 | 0.011 | 0.022 | 0.007 |
| **SalVentAttnB** | 0.008 | 0.004 | 0.009 | 0.067 | 0.000 | 0.037 | 0.007 | 0.007 | 0.029 | 0.118 | 0.020 |
| **SomMotA** | 0.041 | 0.038 | ***0.274*** | 0.023 | 0.014 | 0.002 | 0.001 | 0.000 | 0.000 | 0.000 | 0.030 |
| **SomMotB** | ***0.288*** | 0.041 | 0.003 | 0.080 | 0.001 | 0.000 | 0.017 | 0.002 | 0.010 | 0.001 | 0.017 |
| **TempPar** | *0.156* | 0.015 | 0.000 | 0.017 | 0.003 | 0.020 | 0.094 | 0.000 | 0.043 | 0.028 | 0.024 |
| **VisCent** | 0.002 | 0.004 | 0.001 | 0.005 | 0.027 | 0.000 | ***0.197*** | *0.185* | 0.002 | 0.018 | 0.012 |
| **VisPeri** | 0.000 | 0.051 | 0.000 | 0.019 | 0.001 | 0.004 | 0.001 | ***0.286*** | 0.000 | **0.122** | 0.003 |

***Note:*** *The selected network labels for the respective independent components (ICs) are shown in bold, whereas all cognitive networks that showed a higher Similarity Coefficient than J = 0.15 are shown in italics. No Similarity Coefficient over J = 0.15 was found for IC16 and for IC33.*

**Table S6.** Statistical results for domain-specific network activity

| | expected vs. rest | | unexpected vs. rest | | anomalous vs. rest | | pseudoword vs. rest | | unexpected vs. expected | | anomalous vs. expected | | pseudoword vs. expected | |
|---|---|---|---|---|---|---|---|---|---|---|---|---|---|---|
| | **t-values** | **p-values** | **t-values** | **p-values** | **t-values** | **p-values** | **t-values** | **p-values** | **t-values** | **p-values** | **t-values** | **p-values** | **t-values** | **p-values** |
| **IC04** | 8.548 | **0.000** | 8.498 | **0.000** | 8.006 | **0.000** | 8.565 | **0.000** | -0.891 | 0.576 | 0.655 | 0.790 | 2.126 | 0.114 |
| **IC11** | -4.832 | **0.000** | -4.719 | **0.000** | -5.127 | **0.000** | -6.961 | **0.000** | -0.348 | 0.803 | -1.718 | 0.296 | -3.597 | **0.004** |
| **IC12** | 3.998 | **0.001** | 4.213 | **0.001** | 3.712 | **0.002** | 3.174 | **0.009** | 0.823 | 0.576 | -0.464 | 0.790 | -1.176 | 0.431 |
| **IC13** | 0.756 | 0.556 | 1.671 | 0.192 | 0.761 | 0.453 | 1.320 | 0.216 | -1.377 | 0.490 | -0.078 | 0.998 | 1.059 | 0.431 |
| **IC16** | 1.752 | 0.226 | 1.240 | 0.287 | 2.704 | **0.024** | 1.749 | 0.142 | 1.153 | 0.567 | 1.537 | 0.296 | 0.173 | 0.863 |
| **IC17** | -1.338 | 0.262 | -3.972 | **0.001** | -1.503 | 0.197 | -1.875 | 0.129 | 3.023 | 0.055 | -0.553 | 0.790 | -1.025 | 0.431 |
| **IC18** | -1.584 | 0.226 | -1.187 | 0.287 | -2.227 | 0.061 | -1.583 | 0.151 | -0.819 | 0.576 | -1.570 | 0.296 | -0.191 | 0.863 |
| **IC21** | 0.469 | 0.706 | 0.491 | 0.627 | 0.846 | 0.444 | 0.169 | 0.867 | -0.077 | 0.939 | 0.632 | 0.790 | -0.584 | 0.689 |
| **IC25** | 1.353 | 0.262 | 3.265 | **0.006** | 4.662 | 0.000 | 4.626 | **0.000** | -2.419 | 0.119 | 5.084 | 0.000 | 4.037 | **0.004** |
| **IC33** | 0.096 | 0.924 | 1.146 | 0.287 | 1.110 | 0.337 | 1.667 | 0.145 | -1.766 | 0.320 | 1.826 | 0.296 | 3.615 | **0.004** |
| **IC35** | -1.656 | 0.226 | -1.350 | 0.287 | -1.510 | 0.197 | -2.929 | **0.014** | -0.602 | 0.674 | -0.002 | 0.998 | -1.976 | 0.126 |

***Note:*** *P-values are FDR-corrected at $p < 0.05$. Bold values indicate significant comparisons in the two-sided t-tests.*

**Figure S5**

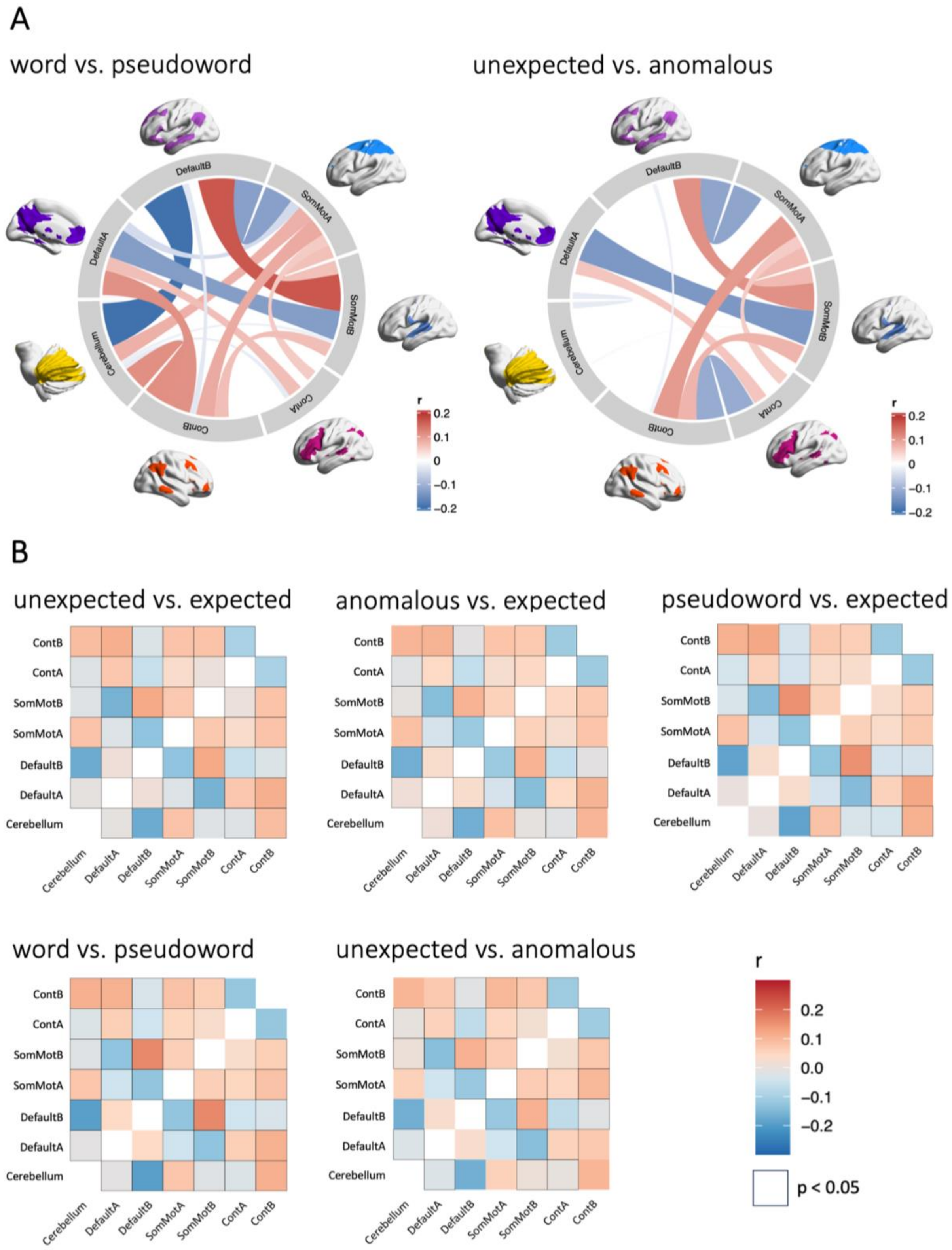

**Figure S5.** *Functional coupling between task-relevant networks. A. Chord diagrams show (additional) significant results of functional coupling between ICA-derived networks for the contrasts "words vs. pseudowords" and "unexpected vs. anomalous". Connectivity values are partial correlations. The color*

*intensity and width of a connection indicate its correlational strength. Higher correlation values (r) indicate positive coupling and negative values indicate decoupling between networks. B. Heatmaps show correlation matrices of functional coupling between ICA-derived networks. The color intensity indicates correlational strength between two networks. Significant correlations are highlighted by black boxes. Higher values indicate positive coupling and negative values indicate decoupling between networks. Abbreviations: ContA/B: Control network A/B; DefaultA/B: Default mode network A/B; SomMotA/B: Somatomotor network A/B.*

**Table S7.** *CPPI connectivity effects on response time contrasting unexpected vs. expected sentence endings*

| | **Log (Reaction times)** | | |
|---|---|---|---|
| ***Predictors*** | *Estimates* | *CI* | *p* |
| **(Intercept)** | 6.72 | 6.41 – 7.03 | **<0.001** |
| **DefaultA__SomMotA** | -0.03 | -0.07 – 0.02 | 0.217 |
| **condition [unexpected]** | -0.09 | -0.10 – -0.07 | **<0.001** |
| **age** | 0.00 | -0.00 – 0.00 | 0.426 |
| **education** | 0.00 | -0.02 – 0.02 | 0.790 |
| **DefaultA__SomMotA × condition [unexpected]** | 0.02 | 0.01 – 0.03 | **0.001** |
| **Random Effects** | | | |
| **$\sigma^2$** | 0.03 | | |
| **$\tau_{00\ subject}$** | 0.01 | | |
| **ICC** | 0.31 | | |
| **$N_{subject}$** | 32 | | |
| **Observations** | 3673 | | |
| **Marginal $R^2$ / Conditional $R^2$** | 0.064 / 0.357 | | |

***Note:*** *Significant effects are marked in bold. Contrasts are treatment coded. P-values were obtained via likelihood ratio tests.* ***Abbreviations:*** *CI: Confidence interval; DefaultA: Default mode network A; SomMotA: Somatomotor network A.*

**Table S8**. *CPPI connectivity effects on response time contrasting anomalous vs. expected sentence endings*

| | Log (Reaction times) | | | Log (Reaction times) | | |
|---|---|---|---|---|---|---|
| ***Predictors*** | *Estimates* | *CI* | *p* | *Estimates* | *CI* | *p* |
| **(Intercept)** | 6.83 | 6.51 – 7.14 | **<0.001** | 6.81 | 6.48 – 7.13 | **<0.001** |
| **SomMotB__ContA** | -0.00 | -0.04 – 0.04 | 0.999 | | | |
| **condition [anomalous]** | -0.23 | -0.24 – -0.22 | **<0.001** | -0.23 | -0.24 – -0.22 | **<0.001** |
| **age** | 0.00 | -0.00 – 0.00 | 0.370 | 0.00 | -0.00 – 0.00 | 0.412 |
| **education** | 0.00 | -0.02 – 0.02 | 0.993 | 0.00 | -0.02 – 0.02 | 0.863 |
| **SomMotB__ContA × condition [anomalous]** | -0.02 | -0.03 – -0.01 | **0.002** | | | |
| **SomMotB__DefaultB** | | | | -0.01 | -0.06 – 0.04 | 0.696 |
| **SomMotB DefaultB × condition [anomalous]** | | | | 0.02 | 0.01 – 0.03 | **0.001** |
| **Random Effects** | | | | | | |
| $\sigma^2$ | 0.03 | | | 0.03 | | |
| $\tau_{00}$ | 0.01 $_{subject}$ | | | 0.01 $_{subject}$ | | |
| **ICC** | 0.31 | | | 0.31 | | |
| **N** | 32 $_{subject}$ | | | 32 $_{subject}$ | | |
| **Observations** | 3654 | | | 3654 | | |
| **Marginal $R^2$ / Conditional $R^2$** | 0.232 / 0.473 | | | 0.233 / 0.473 | | |

***Note:*** *Significant effects are marked in bold. Contrasts are treatment coded. P-values were obtained via likelihood ratio tests.* ***Abbreviations:*** *CI: Confidence interval; ContA: Control network A; DefaultB: Default mode network B; SomMotB: Somatomotor network B.*

#### **Table S9.** *CPPI connectivity effects on response time contrasting pseudoword vs. expected sentence endings*

| | Log (Reaction times) | | | Log (Reaction times) | | | Log (Reaction times) | | | Log (Reaction times) | | | Log (Reaction times) | | | Log (Reaction times) | | |
|---|---|---|---|---|---|---|---|---|---|---|---|---|---|---|---|---|---|---|
| ***Predictors*** | *Estimates* | *CI* | *p* | *Estimates* | *CI* | *p* | *Estimates* | *CI* | *p* | *Estimates* | *CI* | *p* | *Estimates* | *CI* | *p* | *Estimates* | *CI* | *p* |
| **(Intercept)** | 6.78 | 6.48 – 7.08 | **<0.001** | 6.78 | 6.50 – 7.07 | **<0.001** | 6.81 | 6.52 – 7.09 | **<0.001** | 6.76 | 6.45 – 7.06 | **<0.001** | 6.79 | 6.50 – 7.07 | **<0.001** | 6.86 | 6.58 – 7.13 | **<0.001** |
| **DefaultA__ContA** | 0.00 | -0.04 – 0.04 | 0.848 | | | | | | | | | | | | | | | |
| **condition [pseudoword]** | 0.27 | 0.26 – 0.28 | **<0.001** | 0.27 | 0.26 – 0.28 | **<0.001** | 0.27 | 0.26 – 0.28 | **<0.001** | 0.27 | 0.26 – 0.28 | **<0.001** | 0.27 | 0.26 – 0.28 | **<0.001** | 0.27 | 0.26 – 0.28 | **<0.001** |
| **age** | 0.00 | -0.00 – 0.00 | 0.447 | 0.00 | -0.00 – 0.00 | 0.595 | 0.00 | -0.00 – 0.00 | 0.498 | 0.00 | -0.00 – 0.00 | 0.404 | 0.00 | -0.00 – 0.00 | 0.579 | 0.00 | -0.00 – 0.00 | 0.904 |
| **education** | 0.01 | -0.01 – 0.02 | 0.541 | 0.01 | -0.01 – 0.02 | 0.449 | 0.00 | -0.01 – 0.02 | 0.633 | 0.01 | -0.01 – 0.02 | 0.493 | 0.01 | -0.01 – 0.02 | 0.486 | 0.00 | -0.01 – 0.02 | 0.633 |
| **DefaultA__ContA × condition [pseudoword]** | 0.03 | 0.02 – 0.04 | **<0.001** | | | | | | | | | | | | | | | |
| **SomMotA__Cerebellum** | | | | -0.03 | -0.07 – 0.01 | 0.145 | | | | | | | | | | | | |
| **SomMotA__Cerebellum × condition [pseudoword]** | | | | -0.02 | -0.03 – -0.01 | **<0.001** | | | | | | | | | | | | |
| **ContB__DefaultB** | | | | | | | 0.03 | -0.01 – 0.07 | 0.106 | | | | | | | | | |
| **ContB__DefaultB × condition [pseudoword]** | | | | | | | -0.02 | -0.03 – -0.01 | **0.001** | | | | | | | | | |
| **SomMotB__ContB** | | | | | | | | | | -0.02 | -0.06 – 0.03 | 0.442 | | | | | | |
| **SomMotB__ContB × condition [pseudoword]** | | | | | | | | | | 0.01 | 0.01 – 0.02 | **0.003** | | | | | | |
| **SomMotB__DefaultA** | | | | | | | | | | | | | -0.03 | -0.07 – 0.01 | 0.118 | | | |
| **SomMotB__DefaultA × condition [pseudoword]** | | | | | | | | | | | | | 0.01 | 0.00 – 0.02 | **0.007** | | | |
| **SomMotA__DefaultB** | | | | | | | | | | | | | | | | 0.05 | 0.01 – 0.09 | **0.009** |
| **SomMotA__DefaultB × condition [pseudoword]** | | | | | | | | | | | | | | | | -0.01 | -0.02 – -0.00 | **0.021** |
| **Random Effects** | | | | | | | | | | | | | | | | | | |
| **$\sigma^2$** | 0.03 | | | 0.03 | | | 0.03 | | | 0.03 | | | 0.03 | | | 0.03 | | |
| **$\tau_{00}$** | 0.01 $_{subject}$ | | | 0.01 $_{subject}$ | | | 0.01 $_{subject}$ | | | 0.01 $_{subject}$ | | | 0.01 $_{subject}$ | | | 0.01 $_{subject}$ | | |
| **ICC** | 0.31 | | | 0.29 | | | 0.30 | | | 0.31 | | | 0.30 | | | 0.27 | | |
| **N** | 32 $_{subject}$ | | | 32 $_{subject}$ | | | 32 $_{subject}$ | | | 32 $_{subject}$ | | | 32 $_{subject}$ | | | 32 $_{subject}$ | | |
| **Observations** | 5494 | | | 5494 | | | 5494 | | | 5494 | | | 5494 | | | 5494 | | |
| **Marginal $R^2$ / Conditional $R^2$** | 0.279 / 0.505 | | | 0.293 / 0.501 | | | 0.290 / 0.501 | | | 0.278 / 0.502 | | | 0.290 / 0.501 | | | 0.314 / 0.499 | | |

***Note:** Significant effects are marked in bold. Contrasts are treatment coded. P-values were obtained via likelihood ratio tests. **Abbreviations:** CI: Confidence interval; ContA/B: Control network A/B; DefaultA/B: Default mode network A/B; SomMotA/B: Somatomotor network A/B.*
